# Continuous FACS sorting of double emulsion picoreactors with a 3D printed vertical mixer

**DOI:** 10.1101/2025.03.13.643134

**Authors:** Zijian Yang, Samuel Thompson, Yanrong Zhang, Iene Rutten, Julie Van Duyse, Gert Van Isterdael, Lisa Nichols, Jeroen Lammertyn, Hyongsok T Soh, Polly Fordyce

## Abstract

High-throughput screening and directed evolution using microfluidic picoreactors have produced high-activity enzymes. In this approach, a substrate is co-encapsulated with a candidate enzyme and individual picoreactors are sorted based on an activity reporter. While many approaches use water-in-oil droplets (single emulsions) for fluorescence-activated droplet sorting (FADS) on custom-fabricated microfluidic devices that require integrated optics and electronics, recent approaches have lowered the engineering barriers to adoption by using simple microfluidic droplet generators to produce water-in-oil-in-water droplets (double emulsion picoreactors, DEs) that can be sorted with commercial FACS (fluorescence-activated cell sorting). Despite the simplified engineering requirements, high variability in loading rates and low yield during loading are barriers to efficient DE FACS sorting. Here, we optimized surfactants to enhance DE stability and demonstrated that a 3D-printed corkscrew on the sample line acts as a vertical mixer to enable more continuous loading. With these optimized loading conditions, we analyzed 1.17 million DEs in four 10-minute sorting rounds with a mean frequency of 480 Hz (390 Hz including sample exchanges); in a mock sort of 10% fluorescent DEs, we achieved 89.2±1.1% accuracy and 78±0.9% recovery with our optimized loading protocol. Overall, improved ease-of-use and throughput for FACS sortable DEs should expand the accessibility of directed evolution in controlled *in vitro* environments.

## INTRODUCTION

Library selection for directed evolution has produced many feats of macromolecular engineering, particularly in the production of novel enzymes and biocatalysts^1^. Cell-based assays ^2–8^ (e.g. phage display, yeast display, bacterial FACS, etc.) are convenient because they produce protein *in situ* and link genotype and phenotype; however, they impractical for many engineering applications because they are incompatible with extreme reaction conditions (e.g. labile substrates, pH, temperature, toxic compounds, secreted compounds). Microfluidic droplet picoreactors offer a powerful alternative and support reaction conditions incompatible with cell-based screens^9^. These picoreactors physically link a target macromolecule with a functional read-out (e.g. an assay product) through encapsulation, reduce reagent use (1-100 pL per reaction), and are compatible with convenient readouts (e.g. fluorescence^10– 12^, mass^13^, and absorbance^14,15^, or electrochemistry^16^). Microfluidic generators also allow for customizable workflows and for highly controlled mixing and picoinjection.

While single emulsion (water-in-oil) picoreactors have been successfully used for selection via fluorescence-activated droplet sorting (FADS), this approach requires custom microfluidic sorters integrating flow control, optics, and piezoelectrics. To increase accessibility, recent approaches (**Supplementary Table 1**) have developed double emulsion picoreactors (DEs, water-in-oil-in-water) that can be sorted with commercial FACS machines (fluorescence-activated cell sorting). FACS offers multicolor sorting (up to 18 fluorescence channels)^17^ and high peak sorting rates (up to 20 kHz), and FACS sortable DEs have been used for multiple assays including RT PCR^18^, aptamer sensors for enzymatic products^19^, vesicle rupture^20^, and fluorogenic enzyme substrates^9,21–24^. The diameter of sorted particles must be < 30% of the size of the nozzle to prevent DE shearing during sorting^9,19,25–27^ and larger nozzles have slower maximum sort rates^26,28^.

For DEs, a trade-off exists between throughput and control. Heterogenous bulk-generated DEs are generally <20 μm in diameter and can be sorted with smaller, faster nozzles, but low signal-to-noise and high variability in reaction volume size and reagent concentration impede many applications, and applications such as reverse transcription from encapsulated mammalian cells require larger DEs^18,27^. Reflecting a greater desire for control over DE size variation and the composition of reagents in the DE core, the field has shifted to microfluidic DEs (**Figure S1A**) that are monodisperse^26,29–31^ and compatible with mammalian cells^18,27^; however, these DEs tend to be large (30-50 μm) and therefore require 130 μm nozzles to maintain stable breakoff of the sorter stream, which limits peak reported sorting rates to 2-3 kHz (**Figure S1B**).

Despite reported high *peak* sorting rates, maintaining these rates over long time frames is technically challenging. As a result, the vast majority of DE studies in the past decade analyzed <250,000 events and did not evolve a new macromolecule (**Supplementary Table 1, Figure S1C**,**D**). These sorting challenges have been further exacerbated by increasing use of dense fluorocarbon oils compatible with a wide variety of biochemical and cellular reactions, as large dense DEs settle rapidly in the sample tube. As DEs settle, they become inaccessible to the loading tube and the loading rate decays rapidly, limiting throughput until the sample is unloaded and manually resuspended. Additionally, DEs are more fragile than cells because larger particles are subject to more intense shearing forces and DEs lack a cytoskeleton. Sheared DEs frequently clog the FACS sample lines and nozzle, further reducing the overall sorting efficiency^26,28^.

Here, we systematically optimized DE loading rates and stability, with the goal of sorting 1 million DEs per hour (approximately 300 Hz). In this optimization, we varied surfactants and surfactant concentrations and attached a 3D-printed corkscrew onto the FACS loading tubes to prevent DE settling. Using our optimized conditions, we demonstrate the ability to consistently sort at peak loading rates of 500-1000 Hz over several minutes with 89.2±1.1% accuracy and 78±0.9% recovery. With 4 rounds of iterative sample loading, we successfully sorted 10^6^ 40 μm DE picoreactors in <1 hour including sample exchange times. In future work, we anticipate these technical advances will significantly extend the duration of uninterrupted sorting and enable routine analysis of 10^5^-10^6^ member libraries in directed evolution campaigns.

## RESULTS

### Optimizing oil surfactant for double emulsion picoreactor stability

DEs are composed of an inner aqueous layer, a fluorocarbon (e.g. HFE7500) or hydrocarbon (e.g. mineral oil) oil shell, and an outer aqueous layer that allows DEs to be suspended within the FACS sheath fluid (**Figure 1A**). Added surfactants migrate to each water/oil interface, where they lower interfacial surface tension to promote DE formation and prevent coalescence. Some commercially available nonionic droplet surfactants (e.g. dSurf and PicoSurf) have been optimized for improved compatibility with cellular and biochemical assays relative to ionic surfactants and hydrocarbon oils^9,32,33^; however, their impacts on long-term DE stability and sorting remain uncharacterized.

**Figure 1.**
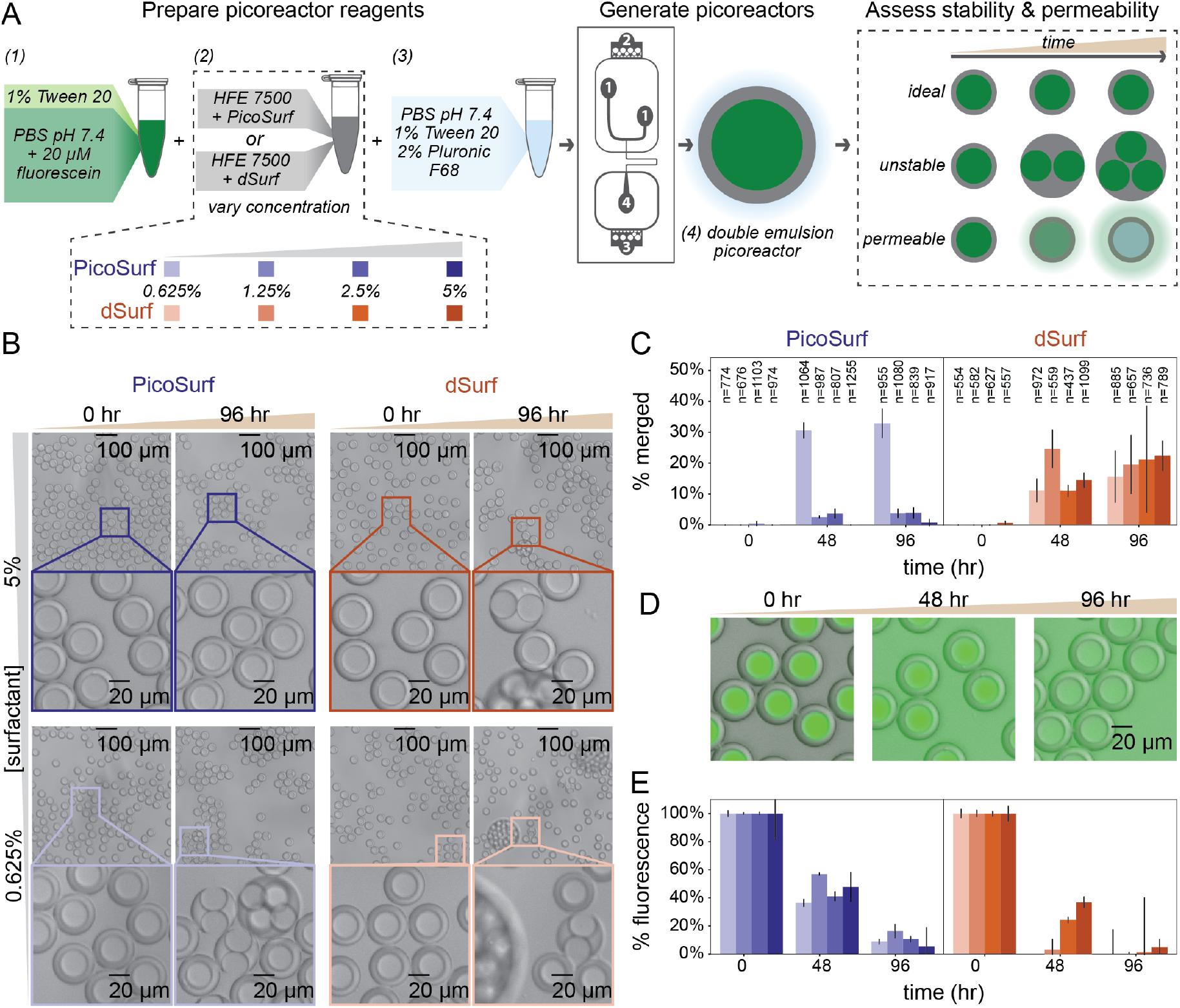
Optimize oil surfactants for double emulsion picoreactor stability. **A)** Workflow for generating DE picoreactors and analyzing stability and permeability by microscopy. Color gradients indicate varying concentrations of PicoSurf (blue) and DSurf (orange); numbered inputs and outputs match ports/channel on the picoreactor generator schematic. **B)** Brightfield microscopy images of picoreactors at 0 and 96 hrs (brown gradient) for 5% and 0.625% surfactant concentrations (grey gradient). Scale bar: 100 μm. Inset images show 5x zoom of (outline) one section of each respective image. Inset scale bar: 20 μm. **C)** Percentage of aqueous cores in merged picoreactors as a function of surfactant condition and time. Error bars represent standard deviation across 3 images; annotations specify the total number of aqueous cores analyzed over all three images. **D)** Merged brightfield and green fluorescence microscopy images for 5% PicoSurf picoreactors at 0, 48, and 96 hrs. Scale bar: 20 μm. **E)** Retained initial fluorescence as a function of surfactant condition and time. Error bars represent standard deviation across three images.

To test the impacts of these surfactants on DE stability, we: (1) generated DEs with HFE7500 oil and 0.625-5% (w/v) of either PicoSurf or dSurf, (2) added 20 μM fluorescein to the inner aqueous phase to label the aqueous core, and (3) monitored DE merging with optical microscopy after 0, 48, and 96 hours (n=437-1255 DE cores per timepoint; **Figures 1B, 1C, S2** and **Supplementary Table 2**). For moderate and higher concentrations of PicoSurf, DEs showed little unwanted merging even after 96 hours (<1% merged for 5% PicoSurf). In contrast, dSurf DEs showed substantial merging that continually increased over time (15-25% merged for all dSurf concentrations after 96 hours and nearly half of the DEs merged in 96 hours for the lowest concentration of PicoSurf). In addition, merged PicoSurf DEs typically contained 2 aqueous cores while dSurf DEs formed large multicore aggregates (**Figure 1B**). Overall, these results indicate that PicoSurf concentrations ≥1.25% are optimal for long-term DE stability.

Surfactant properties can also impact the rate of solute leakage from DEs^34,35^. To test whether conditions that enhance stability alter leakage, we also imaged DEs to quantify the fluorescence of the inner aqueous core during the time course (**Figure 1D**). Over 96 hours, the background-subtracted fluorescein intensity within the inner core decreased to 0-20%. For PicoSurf DEs, the raw fluorescence signal for the DE cores and the background fluorescence asymptotically approached the same value over 96 hours (**Figure S3**), consistent with an equilibration between the core and the outer solution on the order of days. Similar multi-day equilibration kinetics have previously been observed with both single and double emulsions^35,36^. Fluorescein leakage was faster for dSurf DEs, reaching equilibrium around 48 hours. The final fluorescence intensity was generally higher for PicoSurf DEs at the 96-hour timepoint (**Figure 1E, S3**, and **Supplementary Table 3**). Fluorescence leakage was slower for aqueous cores in merged DEs, likely due to a reduced surface area to volume ratio (**Figure S4**). Overall, PicoSurf DEs appear to retain small molecules more effectively than dSurf.

### PicoSurf-containing picoreactors exhibit more consistent sorting rates

Successfully characterizing large DE libraries requires not only that DEs remain stable but also that they can be loaded into the FACS machine at consistent rates over time. To test for impacts of surfactant conditions on DE loading and fluorescence retention, we used our previously generated double emulsions (**Figure 1**) with 20 μM fluorescein and 1.25%, 2.5%, or 5% of either PicoSurf or dSurf, incubated for 96 hours to match the imaging in **Figure 1**, aliquoted 50 μL (65,000-75,000 DEs) of each sample into a small volume library tube, loaded samples onto a BD FACS Aria III, and quantified monocore purity and loading rate over time (**Figure 2A**); 3 replicates of each sample were loaded under 300 rpm agitation for 90 seconds. For many samples, DE events did not appear for the first 1-2 minutes of loading; in these cases, we started recording data after event rates first reached 20-40 events/sec (**Figure 2B, Figures S5**,**6**). Overall, PicoSurf DEs exhibited a more consistent loading pattern: loading rates increased over 10-20 seconds, peaked between 500-2000 Hz, and quickly dropped below 300 Hz (often dropping below 100 Hz by the end of the run) (**Figure 2B**). In contrast, dSurf DEs exhibited multiple brief loading rate spikes and frequent periods below 100 Hz (**Figure 2B**). The median total monocore DE yield after 90 seconds of loading was significantly higher for PicoSurf samples relative to dSurf, with 5% PicoSurf DEs yielding a median of 33882±21643 DEs over 90 seconds (equivalent to a constant integrated sorting rate of 375 Hz) (**Supplementary Table 4**). Excess sample remained in all cases such that total yield is not comparable to total generated DEs. Consistent with the high stability we observed for PicoSurf DEs from microscopy analysis (**Figure 1**), monocore purity was higher for PicoSurf (**Figure 2C, Supplementary Table 5**) than for dSurf and exceeded 95%. To analyze DE permeability, we analyzed the fluorescein fluorescence intensity (488/525 nm laser excitation and emission). PicoSurf DEs had higher mean fluorescence than dSurf DEs (**Figure 2D, Supplementary Table 6**), with the trends in intensities observed by FACS mirroring those observed by microscopy (**Figures 2D, 1E**). For the range of surfactant concentrations analyzed both by FACS and microscopy, the highest fluorescence signal for dSurf DEs was equivalent to the lowest fluorescence signal for PicoSurf DEs. Overall, PicoSurf DEs demonstrate better loading, stability, and fluorophore retention than dSurf reactors.

**Figure 2.**
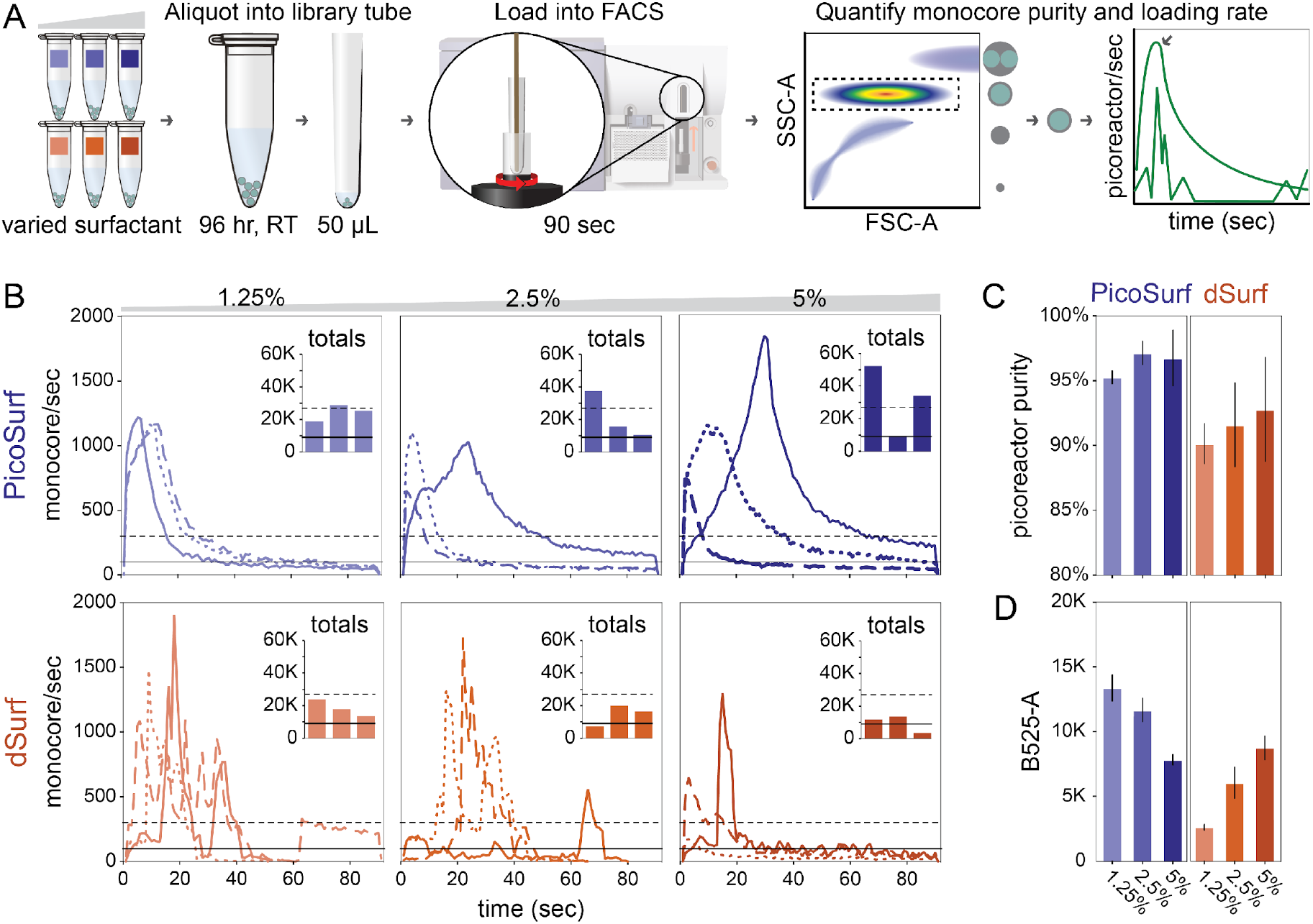
FACS analysis to optimize oil surfactant for picoreactor loading. **A)** Workflow for analyzing monocore DE picoreactor FACS loading rates as a function of surfactant condition. Example diagram (far right) depicts inconsistent and consistent (arrow) loading. **B)** Monocore DE picoreactor loading rates over three independent 90 sec runs (solid line, long dashes, short dashes). Data are colored by surfactant condition as in **Figure 1**. Black lines indicate benchmark rates: 100 (solid) or 300 (dashed) picoreactors/sec. Inset bar charts indicate total monocore picoreactors accumulated for each run. **C)** Picoreactor purity (percentage of monocore picoreactors) out of all FACS scattering events. Error bars represent standard deviation over 3 independent runs. **D)** Monocore picoreactor permeability as mean ratio of green fluorescence (B525) over FSC-A. Error bars represent standard deviation over three independent runs.

### Optimizing picoreactor resuspension for consistent FACS loading using a corkscrew vertical mixer

Sorting 10^6^ DEs in one hour requires a mean sorting frequency of approximately 300 Hz. Ideally, this includes both setup (sample loading, cleaning, etc.) and sorting time. Because intermittent manual sample agitation is recommended even for less dense samples (e.g. cells), we predicted that DE settling caused the observed steep declines in sorting rates. To identify parameters that could impact settling velocity, we modeled DEs using Stokes law (**Figure S7**). Changing physical parameters of the DEs (e.g. oil density, outer solution viscosity) can reduce settling; however, these changes have the potential to impact DE stability and/or the encapsulated assay. As an alternate approach, we explored if improving mixing by increasing agitation could maintain DEs in suspension and enable 300+ Hz sorting over longer durations (e.g. larger sample volumes). To test impacts of varying sample volumes, we loaded samples using either small-volume library tubes or standard FACS tubes; to test impacts of improved agitation, we designed a custom 3D printed corkscrew and attached it to the FACS sample line inserted into a standard FACS tube (**Figure 3A)**. As the corkscrew is wider than the library tubes (width: 8.8 mm, volume: 1.2 mL), this was only tested in standard FACS tubes (width: 12 mm, volume: 5 mL). The corkscrew’s unique design enhances sample agitation by creating a vertical mixing action. Its right-turning (clockwise) thread creates an upward current in solution that counteracts gravity when rotational agitation is provided by the sorter sample loader. We expected this upward current to reduce the DEs settling. Specifically, we tested 3 conditions to examine this prediction. For each condition, we loaded three 600 μL aliquots of 5% PicoSurf DEs (approximately 10 min of DE generation or 750,000-850,000 DEs) and sorted for at least 5 minutes. Peak rates using library tubes and the larger sample volume ranged from 300-2000 Hz with total DE yields of 50,000-175,000 (**Figure 3B, Figure S8, Supplementary Table 7**); as with prior sorts, excess sample remained in all cases. Peak rates with FACS tubes ranged from 300-1500 Hz but sorting rates and total DE yield were consistently lower than with library tubes (yield ranged from 20,000-120,000 DEs) (**Figure 3C, Figure S9, Supplementary Table 8**). Peak rates with the corkscrew and FACS tubes ranged between 1000-1500 Hz and were more continuous than in conditions without the corkscrew, making it possible to extend the sort time to 10 minutes (**Figure 3D, Figure S10, Supplementary Table 9**). Total DE yields for the longer sort times ranged 270,000-325,000 and were more consistent than in conditions without the corkscrew. The FACS tube alone decreased loading rates, but adding the corkscrew resulted in more reliable peak yields and more continuous loading rates.

**Figure 3.**
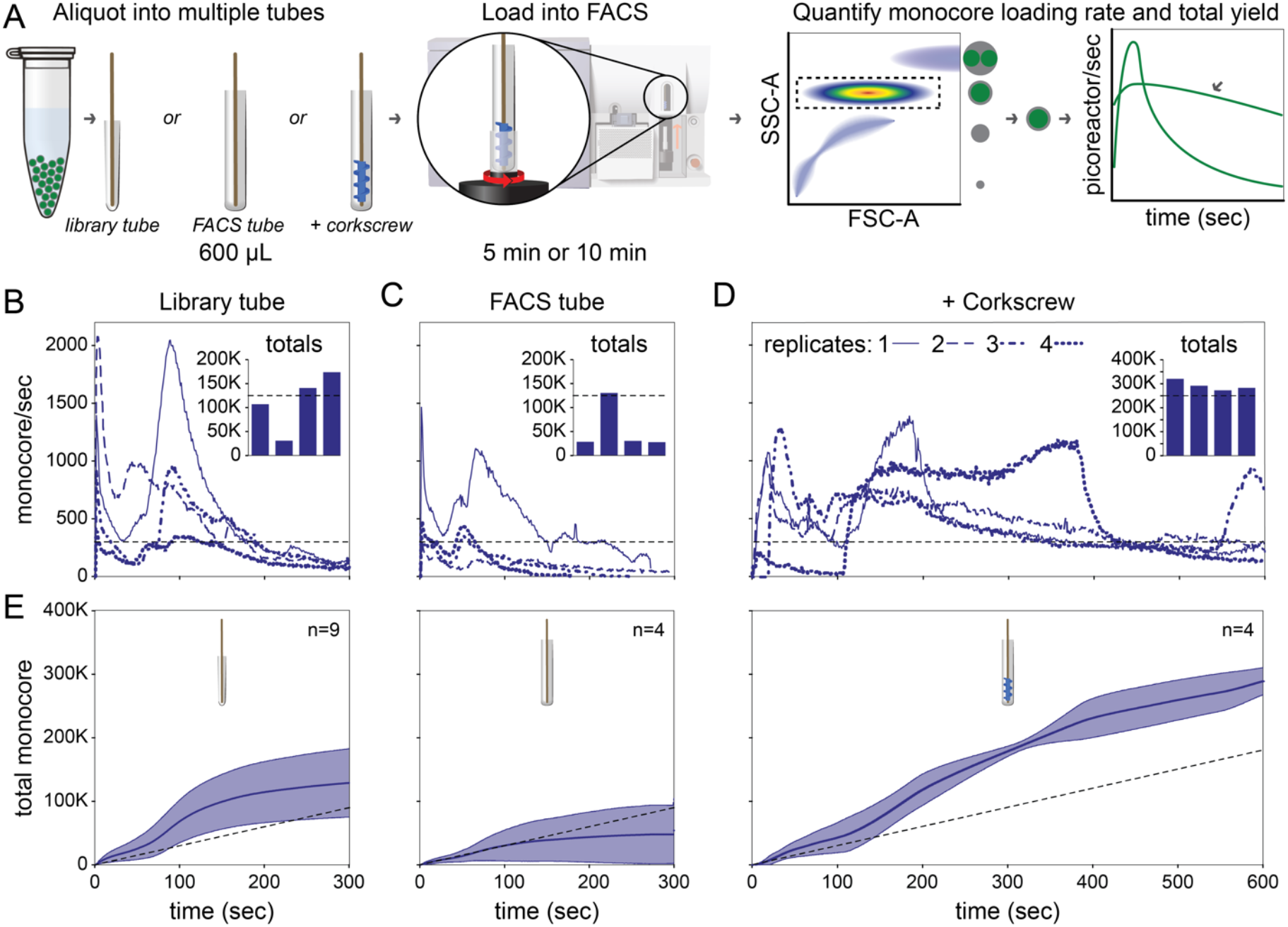
FACS analysis to increase sample agitation for improved picoreactor loading. **A)** Workflow for analyzing 5% Picosurf monocore DE picoreactor FACS loading rates across sample tubes. Example diagram (far right) depicts unsustained and continuous (arrow) loading. **B-D)** Monocore picoreactor loading rates over four independent runs (solid line, long dashes, short dashes, alternating dashes). Runtimes are 5 minutes in B (library tube) and C (FACS tube) or 10 minutes in D (FACS tube with corkscrew). Dashed black lines indicate benchmark rate of 300 picoreactors/sec. Inset bar charts indicate total monocore picoreactors accumulated for each run. **E)** Mean total accumulated picoreactors as a function of run time. n-values indicate number of independent FACS runs for each loading condition. Error bars represent standard deviation over all n runs; dashed black lines indicate total picoreactors at benchmark rate of 300 picoreactors/sec.

We next examined which loading conditions would meet our benchmark of sorting 10^6^ DEs in one hour (equivalent to sustained 300 Hz loading, **Figure 3E**). Both the library tube and FACS tube + corkscrew conditions had mean sorting rates that were consistently higher than 300 Hz. With library tubes, we analyzed 1.16 x 10^6^ DEs over nine 5-minute runs; using FACS tubes with the corkscrew, we analyzed 1.17 x 10^6^ DEs over four 10-minute runs, equivalent to a mean frequency of 480 Hz. Even allowing for 2 minutes of sample handling during each sample exchanges, this would maintain a mean frequency of 390 Hz. Mean sorting rates noticeably decrease after 5 minutes, likely from either sample settling or sample depletion. These results demonstrate that surfactant optimization results in an efficient workflow for sorting double emulsion reactors and that mechanically resuspending the pellet of dense DEs allows for more continuous sorting protocols.

### FACS sorting with corkscrew-assisted loading preserves picoreactor integrity

Screening libraries in DEs further requires accurate recovery of desired populations via FACS sorting (without contamination from unwanted DEs). To demonstrate that the addition of the corkscrew did not compromise DE integrity or sorting, we attempted to select only fluorescent DEs from a population of 10:1 blank-to-fluorescence DEs (approximately 660 μL) using the FACS tube + corkscrew loading condition (**Figure 4A**). To ensure robust linkage between detected and sorted events, we first manually calibrated the FACS drop delay by counting the recovery of 50 DEs sorted onto a glass slide, yielding a recovery rate of 78±0.9% at the optimal delay setting of 22.32 cycles (**Figure 4B**). Using this optimal delay setting, we then detected 354,041 FACS events (**Figure 4C, S11**) including 303,833 monocore DEs (86.9%); consistent with the expected loaded ratio, 11.2% of monocore DEs were fluorescent (MFI > 10^3^; **Figure 4D**).

**Figure 4.**
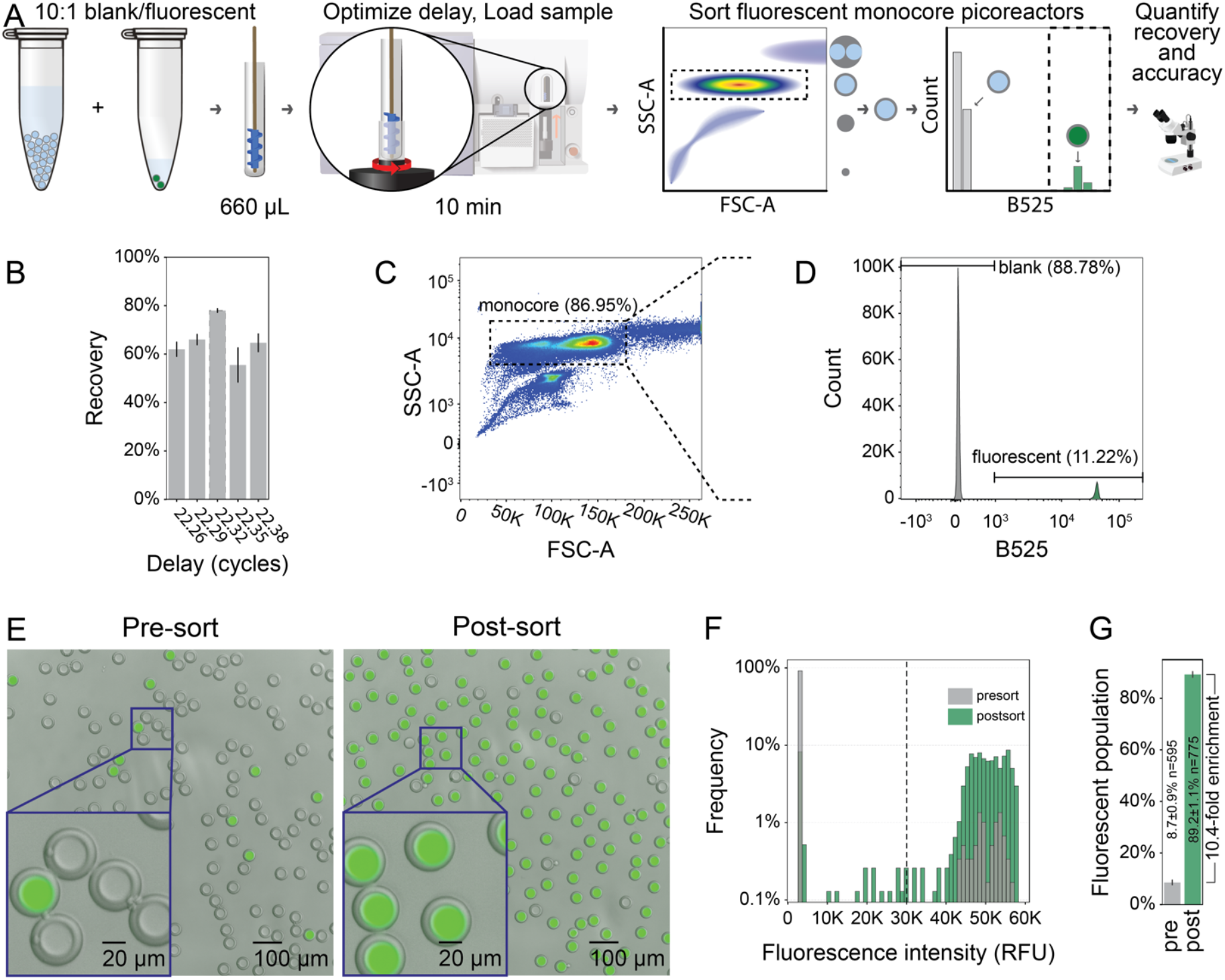
Model FACS sorting to validate optimized double emulsion generation and loading. **A)** Workflow for FACS sorting of rare fluorescent monocore double emulsion picoreactors. **B)** Mean picoreactor recovery as a function of droplet delay. Error bars represent standard deviation over three measurements. **C)** FSC-A vs. SSC-A plot from FACS sorting; dashed line indicates monocore picoreactor gate. **D)** Green fluorescence (B525) intensity histogram for all monocore picoreactors; picoreactors with intensity >10^3^ were gated as fluorescent. **E)** Merged brightfield and green fluorescence images for picoreactors preand post-sort. **F)** Quantification of picoreactor fluorescence from three preand post-sort images. Picoreactors with an intensity >30,000 RFUs (dashed line) were consider fluorescently labeled. **G)** Mean percentage of fluorescent picoreactors in preand post-sort images. Error bars represent the standard deviation of percentages over three images.

To test the ability to successfully recover only desired DEs, we sorted 30,000 fluorescent DEs and analyzed the preand post-sort populations with fluorescence microscopy (**Figure 4E**). While the unsorted sample remaining in the FACS tube showed signs of mechanical damage (**Figure S12**), the sorted population consisted of predominantly intact monocore DEs. Before and after sorting, 8.7±0.9% (n=595) and 89.2±1.1% (n=775) of DEs were fluorescent, respectfully. This selection represented a 10.4-fold enrichment (**Figure 4F**). These results demonstrate that DEs loaded with increased agitation can be efficiently sorted by FACS and recovered intact.

## DISCUSSION

No published papers using the relatively large and monodisperse DEs preferred for modern molecular and cell biology applications have reported analyzing more than 250K DEs total – with one exception where DEs were osmotically shrunk before sorting^29^. This limit likely stems from technical challenges associated with FACS loading that prevent consistent sorting of large DE libraries. To address this, we systematically examined how different surfactant conditions and sample loading methods impact FACS sorting throughput for large (40 μm) microfluidic DEs. DEs with 5% PicoSurf were stable for multiple days and could be consistently sorted via FACS machine over 10-minute runs when loaded using a 3D-printed corkscrew and standard FACS tubes. Using these optimized conditions, we analyzed 1.17 million DEs in four 10-minute sorting runs with sustained sorting rates of 500-1000 Hz and 89.2±1.1% sort accuracy, laying the foundation for future work screening large DE libraries.

Parallelization over multiple FACS sorters (standard for single cell workflows) is an obvious next step for increasing throughput. As an alternative, further optimization of DE size and loading methods could enable sorting of 10^7^-10^8^ large, monodisperse DEs per day using a single sorter. Published work has demonstrated generation of 15 μm^24^ and 20-23 μm^19,29^ monodisperse DEs; as maximum sort rate is inversely correlated with DE size, these >22 μm DEs have been sorted at peak rates of 6-8 Hz. For assays compatible with DEs of this size, maintaining these higher peak rates would boost throughput an additional 20-fold relative to this work. In addition, the incorporation of other mixer geometries (*e*.*g*. vertical ribbon mixers that produce poloidal mixing currents^37^) or the use of density matched oils^38^ or outer fluids^39^ could maintain DE suspension for longer times, boosting ease of sorting by increasing the duration of hands-off sort times.

Optimizing high-throughput sorting of even larger DEs (50-500 μm) on commercial sorters^40–42^ could open new selection assays for organoids, filamentous fungi, and bacterial communities and enable lower signal-to-noise detection methods such as absorption or morphology. Continual end-to-end optimization of FACS sortable DEs will develop this technology into a modular plug-and-play platform that is compatible with a diverse range of assays and encapsulated inputs.

## METHODS

Extended methods are available in Supplementary Information and on the Open Science Framework (OSF) at https://osf.io/jzyve/.

### Picoreactor generation

Double emulsion picoreactors were generated from custom fabricated PDMS devices using previously reported designs^9,18,25,27,30,31^. All input solutions were loaded into plastic syringes (BD) and connected to the PDMS device via LDPE medical tubing (Scientific Commodities Inc.). Syringe pumps drove the flow of reagents into the device, with standard flow rates of 2x25 μL/hr for two identical inner solutions of PBS (Gibco) with 1% (w/v) Tween-20 (Fisher), 125 μL/hr for the picoreactor fluorocarbon oil with the indicated concentration of dSurf (Fluigent) or PicoSurf (SphereFluidics), and 3500-3700 μl/hr for the outer sheath solution of PBS with 1% (w/v) Tween-20 and 2% (w/v) Pluronic F68 (Millipore Sigma). The PBS inner solution was labeled with 20 μM Fluorescein (Fluka).

### Microscopy

Picoreactors were imaged with bright field and fluorescence microscopy (Nikon). Flatfield correction, particle detection, and fluorescence quantification were performed using custom python scripts and the cv2/OpenCV library.

### FACS

Picoreactors were sorted on a FACSAria II cell sorter (BD) with a 130 μm nozzle. Laser delays were manually calibrated with 32 μm AccuCount Ultra Rainbow beads (Spherotech). Forward and side scattering voltage were manually optimized using DE samples; droplet delays were manually calibrated by sorting 50 DEs onto a glass slide at each setting and manually counting recovered picoreactors by optical microscopy. Detector voltages were optimized using DE samples labeled with fluorescein. Samples were loaded in the volume and configuration (library tube, FACS tube, FACS tube + corkscrew) as described in the figures.

## Supporting information

Supplementary Table 1 - Literature

## ASSOCIATED CONTENT

### Supporting Information

The Supporting Information is available free of charge on the ACS Publications website. Additional images, CSV datafiles, FACS workspace files, and Autocad files are available in the Open Science Framework (OSF) at https://osf.io/jzyve/. Stereolithography files for the 3D corkscrew are available on Thingiverse at https://www.thingiverse.com/thing:6675740.

## AUTHOR INFORMATION

### Author Contributions

ZY and SMT conceived the project. ZY and SMT prepared all experimental samples. IR, JVD, and GVI designed and prepared the corkscrew mixer. SMT performed all microscopy. ZY, YZ, and LN performed all FACS analysis and sorting. PF, HTS, JL, and LN provided supervision. ZY, SMT, and PF prepared all figures and drafted the manuscript. SMT, PF, and JL provided funding. All authors provided manuscript edits and have given approval to the final version of the manuscript. ‡These authors contributed equally: ZY and SMT.

## ACKNOWLEDGMENT

The authors would like to thank the members of the Fordyce lab for helpful discussion and insightful feedback and the VIB-IRC workshop for help with the design and 3D printing of the corkscrew mixer. This work was supported by a Garden Grant from the Homeworld Collective and NIH DP1CA290563 awarded to PMF. Data was collected on an instrument in the Shared FACS Facility obtained using NIH S10 Shared Instrument Grants S10RR027431-01 and S10RR025518-01. SMT is supported by the Arnold and Mabel Beckman Foundation through the Arnold O. Beckman Postdoctoral Fellowship in Chemical Sciences. ZY was supported by the Sanjiv Sam Gambhir - Philips Fellowship in Precision Health Program in the Department of Radiology of Stanford University. PMF is a Chan Zuckerberg Biohub Investigator.

## ABBREVIATIONS

DE: double emulsion or double emulsion picoreactor
FACS: fluorescence-activated cell sorting;
FADS: fluorescence-activated droplet sorting
MFI: mean fluorescence intensity;
RFU: relative fluorescence unit;
RT PCR: real-time polymerase chain reaction.

---

Insert Table of Contents artwork here

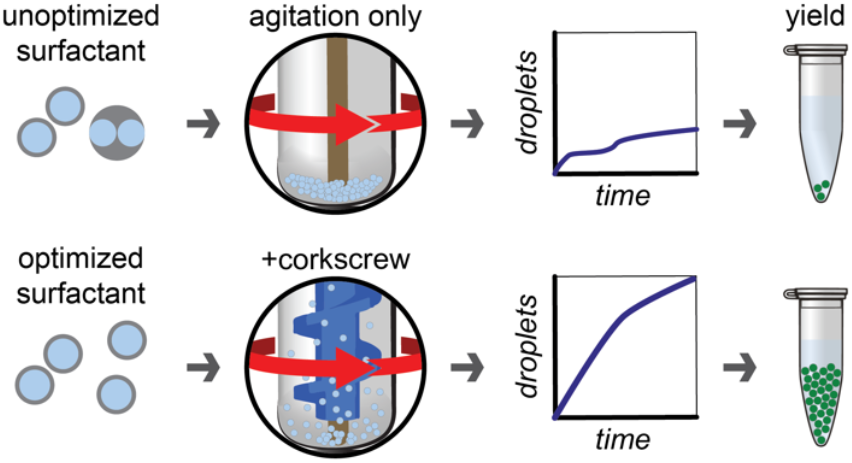

