## Supplementary Information for "Continuous FACS sorting of double emulsion picoreactors with a 3D printed vertical mixer"

**Affiliations**
1) Department of Radiology, Stanford University, Stanford, CA
2) Department of Genetics, Stanford University, Stanford, CA
3) Department of Bioengineering, Stanford University, Stanford, CA
4) Stanford Shared FACS Facility, Stanford University, Stanford, CA
5) Department of Biosystems - Biosensors group, KU Leuven, Leuven, Belgium
6) VIB Flow Core, VIB Technologies, Ghent, Belgium
7) VIB Center for Inflammation Research, Department of Biomedical Molecular Biology, Ghent University, Ghent, Belgium
8) Center for Molecular and Genetic Medicine, Stanford University, Stanford, CA
9) Department of Electrical Engineering, Stanford University, Stanford, CA
10) ChEM-H Institute, Stanford University, Stanford, CA
11) Chan Zuckerberg Biohub, San Francisco, CA

**Supplementary Files**
Supplementary files are separate files available at https://osf.io/jzyve/. All data are paired with any custom python scripts used for analysis. Data Supplements 2, 4, and 6 include workspace files for the FlowJo commercial software suite.

**Data Supplement 1** is microscopy data (.tif) used for analysis in Figure 1
**Data Supplement 2** is FACS workspace files (.fcs) used for analysis in Figure 2.
**Data Supplement 3** is real-time loading rates (.csv) used for analysis in Figure 2.
**Data Supplement 4** is FACS workspace files (.fcs) used for analysis in Figure 3.
**Data Supplement 5** is real-time loading rates (.csv) used for analysis in Figure 3.
**Data Supplement 6** is a FACS workspace file (.fcs) used for analysis in Figure 4.
**Data Supplement 7** is microscopy data (.tif) used for analysis in Figure 4.

**Supplementary Tables**
Supplementary Tables 1 is included as a separate .csv file available at https://osf.io/jzyve/.
Supplementary Tables 2-9 are included below

**Supplementary Table 1** compiles literature data on double emulsion FACS sorting.
**Supplementary Table 2** contains source data for double emulsion stability (via microscopy) from Figure 1C.
**Supplementary Table 3** contains source data for double emulsion leakage (via microscopy) from Figure 1D.

**Supplementary Table 4** contains source data for total yields from Figure 2B.
**Supplementary Table 5** contains source data for double emulsion stability (via FACS) from Figure 2C.
**Supplementary Table 6** contains source data for double emulsion leakage (via FACS) from Figure 2D.
**Supplementary Table 7** contains source data for total yields in Figures 3B and S6 (library tube).
**Supplementary Table 8** contains source data for total yields in Figures 3C and S7 (FACS tube).
**Supplementary Table 9** contains source data for total yields in Figures 3D and S8 (FACS tube + corkscrew).

**Supplementary Figures**


**Supplementary Figure 1:** ***Analysis of published FACS sortable double emulsion generation methods and throughput.***
**Supplementary Figure 2:** ***Automated detection of double emulsions.***
**Supplementary Figure 3:** ***Raw fluorescence and background in fluorescein containing double emulsion picoreactors.***
**Supplementary Figure 4:** ***Merged multicore double emulsion picoreactors show higher retention of fluorophore in the aqueous cores.***
**Supplementary Figure 5:** ***FACS analysis of double emulsion picoreactor loading and uniformity with PicoSurf surfactant.***
**Supplementary Figure 6:** ***FACS analysis of double emulsion picoreactor loading and uniformity with dSurf surfactant.***
**Supplementary Figure 7:** ***Stokes law model of double emulsion picoreactor settling.***
**Supplementary Figure 8:** ***FACS analysis of double emulsion picoreactor loading and uniformity with library tube loading.***
**Supplementary Figure 9:** ***FACS analysis of double emulsion picoreactor loading and uniformity with FACS tube loading.***
**Supplementary Figure 10:** ***FACS analysis of double emulsion picoreactor loading and uniformity with corkscrew mixer loading.***
**Supplementary Figure 11:** ***FACS analysis of double emulsion picoreactor loading and uniformity during rare population selection with corkscrew mixer loading.***
**Supplementary Figure 12:** ***Sample agitation with corkscrew mixer results in merged double emulsion picoreactors.***


**Extended Methods**


**Operating microfluidic devices**
*Preparing stock reagents*
*PDMS device generation*
*Generating double emulsion picoreactors*
**Imaging double emulsion picoreactors**
**FACS sorting microfluidics picoreactors**
*3D printing a vertical mixer*
*FACS calibration, analysis, and sorting*

**Supplementary Tables**

**Supplementary Table 2.** ***Double emulsion picoreactor stability***

| Surfactant | Concentration (%) | Time (hr) | % Merged | Std. Dev. |
|---|---|---|---|---|
| PicoSurf | 5 | 0 | 0 | 0 |
| PicoSurf | 5 | 48 | 0 | 0 |
| PicoSurf | 5 | 96 | 0.8 | 1.1 |
| PicoSurf | 2.5 | 0 | 0 | 0 |
| PicoSurf | 2.5 | 48 | 3.7 | 3.8 |
| PicoSurf | 2.5 | 96 | 1.6 | 1.8 |
| PicoSurf | 1.25 | 0 | 0.5 | 0.7 |
| PicoSurf | 1.25 | 48 | 2.6 | 0.4 |
| PicoSurf | 1.25 | 96 | 3.8 | 1.4 |
| PicoSurf | 0.625 | 0 | 0 | 0 |
| PicoSurf | 0.625 | 48 | 30.7 | 2.5 |
| PicoSurf | 0.625 | 96 | 33.0 | 4.7 |
| dSurf | 5 | 0 | 0.7 | 0.5 |
| dSurf | 5 | 48 | 14.4 | 2.4 |
| dSurf | 5 | 96 | 22.4 | 4.8 |
| dSurf | 2.5 | 0 | 0 | 0 |
| dSurf | 2.5 | 48 | 11.0 | 1.9 |
| dSurf | 2.5 | 96 | 21.2 | 17.2 |
| dSurf | 1.25 | 0 | 0 | 0 |
| dSurf | 1.25 | 48 | 24.7 | 6.1 |
| dSurf | 1.25 | 96 | 19.6 | 9.5 |
| dSurf | 0.625 | 0 | 0 | 0 |
| dSurf | 0.625 | 48 | 11.2 | 3.8 |
| dSurf | 0.625 | 96 | 15.6 | 8.4 |

**Supplementary Table 3.** ***Double emulsion picoreactor leakage***

| Surfactant | Concentration (%) | Time (hr) | Raw | Std. Dev. | Background | Std. Dev | % Initial normalized fluorescence | Std. Dev. |
|---|---|---|---|---|---|---|---|---|
| PicoSurf | 5 | 0 | 4503282 | 394470 | 369347 | 7131 | 100 | 18.8 |
| PicoSurf | 5 | 48 | 3705493 | 42014 | 1733258 | 20242 | 47.7 | 10.5 |
| PicoSurf | 5 | 96 | 1980220 | 7699 | 1751890 | 10797 | 5.5 | 13.5 |
| PicoSurf | 2.5 | 0 | 5251892 | 41296 | 425799 | 33731 | 100 | 1.1 |
| PicoSurf | 2.5 | 48 | 3668186 | 88533 | 1687687 | 33859 | 41.0 | 3.4 |
| PicoSurf | 2.5 | 96 | 2183751 | 96607 | 1674179 | 90886 | 10.6 | 2.0 |
| PicoSurf | 1.25 | 0 | 5351659 | 6079 | 540333 | 18126 | 100 | 0.6 |
| PicoSurf | 1.25 | 48 | 4292391 | 23995 | 1553826 | 36156 | 56.8 | 1.0 |
| PicoSurf | 1.25 | 96 | 2489995 | 45378 | 1704713 | 19200 | 16.3 | 4.8 |
| PicoSurf | 0.625 | 0 | 5450156 | 39020 | 530634 | 23003 | 100 | 2.2 |
| PicoSurf | 0.625 | 48 | 3539582 | 84865 | 1749421 | 62706 | 36.4 | 2.6 |
| PicoSurf | 0.625 | 96 | 2100859 | 20312 | 1670076 | 22014 | 8.7 | 1.7 |
| dSurf | 5 | 0 | 5760462 | 154810 | 866000 | 271076 | 100 | 5.4 |
| dSurf | 5 | 48 | 3472621 | 15577 | 1661506 | 25407 | 37.0 | 3.8 |
| dSurf | 5 | 96 | 1901899 | 52795 | 1664633 | 55762 | 4.8 | 5.6 |
| dSurf | 2.5 | 0 | 5677194 | 37768 | 593605 | 11011 | 100 | 1.7 |
| dSurf | 2.5 | 48 | 2909309 | 24959 | 1680691 | 8835 | 24.2 | 2.2 |
| dSurf | 2.5 | 96 | 1838817 | 12102 | 1774988 | 19312 | 1.2 | 39.0 |
| dSurf | 1.25 | 0 | 4786802 | 88548 | 419831 | 46466 | 100 | 2.5 |
| dSurf | 1.25 | 48 | 1804835 | 57202 | 1669226 | 55489 | 3.1 | 7.4 |
| dSurf | 1.25 | 96 | 1529198 | 70717 | 1707714 | 77364 | -4.1 | -5.5 |
| dSurf | 0.625 | 0 | 4870482 | 94764 | 403754 | 25391 | 100 | 3.1 |
| dSurf | 0.625 | 48 | 1629809 | 50530 | 1721893 | 48272 | -2.1 | -0.9 |
| dSurf | 0.625 | 96 | 1620654 | 59529 | 1772440 | 41539 | -3.4 | -20.8 |

**Supplementary Table 5.** ***Monocore Double emulsion picoreactor yield from FACS – Surfactant screen***

| Surfactant | Concentration (%) | Sort time (sec) | Gate 1 (% passing) | Gate 2 (% passing) | Gate 3 (% passing) | Final |
|---|---|---|---|---|---|---|
| PicoSurf | 5 | 90 | 34945 (98.9%) | 24547 (99.3%) | 34293 (98.8%) | 33882 |
| PicoSurf | 5 | 90 | 9345 (99.6%) | 9306 (99.6%) | 9267 (99.6%) | 9229 |
| PicoSurf | 5 | 90 | 55415 (98.0%) | 54331 (98.6%) | 53556 (97.8) | 52369 |
| PicoSurf | 2.5 | 90 | 10770 (98.2%) | 10573 (99.8%) | 10547 (99.8%) | 10527 |
| PicoSurf | 2.5 | 90 | 16182 (98.4%) | 15919 (99.7%) | 15867 (99.5%) | 15794 |
| PicoSurf | 2.5 | 90 | 39116 (97.3%) | 38047 (99.3%) | 37795 (99.4%) | 37588 |
| PicoSurf | 1.25 | 90 | 26551 (98.1%) | 26039 (97.6%) | 25415 (99.7%) | 25337 |
| PicoSurf | 1.25 | 90 | 30357 (98.1%) | 29769 (96.9%) | 28861 (99.6%) | 28752 |
| PicoSurf | 1.25 | 90 | 19632 (98.3%) | 19297 (97.6%) | 18826 (99.8%) | 18786 |
| dSurf | 5 | 90 | 3505 (97.3%) | 3410 (99.7%) | 3401 (100.0%) | 3400 |
| dSurf | 5 | 90 | 15276 (91.3%) | 13950 (98.5%) | 13745 (98.9%) | 13600 |
| dSurf | 5 | 90 | 12860 (93.6%) | 12044 (99.2%) | 11945 (99.4%) | 11873 |
| dSurf | 2.5 | 90 | 18522 (91.4%) | 16925 (98.4%) | 16661 (98.4%) | 16399 |
| dSurf | 2.5 | 90 | 21713 (93.4%) | 20283 (98.9%) | 20053 (98.8%) | 19814 |
| dSurf | 2.5 | 90 | 7645 (96.2%) | 7351 (99.2%) | 7291 (99.6%) | 7262 |
| dSurf | 1.25 | 90 | 14989 (95.6%) | 14328 (97.2%) | 13930 (97.1%) | 13525 |
| dSurf | 1.25 | 90 | 19889 (94.3%) | 18755 (96.6%) | 18119 (97.2%) | 17613 |
| dSurf | 1.25 | 90 | 25960 (95.8%) | 24885 (97.8%) | 24341 (97.7%) | 23784 |

**Supplementary Table 6.** ***Monocore Double emulsion picoreactor purity after 96 hours from FACS***

| Surfactant | Concentration (%) | Time (hr) | % Monocore | Std. Dev. |
|---|---|---|---|---|
| PicoSurf | 1.25 | 96 | 95.27 | 0.51 |
| PicoSurf | 2.5 | 96 | 97.15 | 0.91 |
| PicoSurf | 5 | 96 | 96.74 | 2.14 |
| dSurf | 1.25 | 96 | 90.1 | 1.53 |
| dSurf | 2.5 | 96 | 91.6 | 3.24 |
| dSurf | 5 | 96 | 92.8 | 4.01 |

**Supplementary Table 7.** ***Estimated fluorophore retention after 96 hours from FACS***

| Surfactant | Concentration (%) | Time (hr) | B525-A/FSC-A | Std. Dev. |
|---|---|---|---|---|
| PicoSurf | 1.25 | 96 | 0.133 | 0.0108 |
| PicoSurf | 2.5 | 96 | 0.117 | 0.0090 |
| PicoSurf | 5 | 96 | 0.0808 | 0.0040 |
| dSurf | 1.25 | 96 | 0.0302 | 0.0063 |
| dSurf | 2.5 | 96 | 0.0606 | 0.0120 |
| dSurf | 5 | 96 | 0.0328 | 0.0414 |

**Supplementary Table 9.** ***Monocore double emulsion picoreactor yield from FACS – Library tubes***

| Surfactant | Concentration (%) | Sort time (sec) | Gate 1 (% passing) | Gate 2 (% passing) | Gate 3 (% passing) | Final |
|---|---|---|---|---|---|---|
| PicoSurf | 5 | 300 | 178668 (97.8%) | 174653 (99.5%) | 1738642 (99.9%) | 173744 |
| PicoSurf | 5 | 300 | 14683 (98.6%) | 14730 (99.8%) | 144402 (100.0%) | 144341 |
| PicoSurf | 5 | 300 | 157210 (98.0%) | 154001 (99.6%) | 153399 (99.9%) | 153315 |
| PicoSurf | 5 | 300 | 57049 (98.9%) | 56424 (99.7%) | 56240 (99.8%) | 56132 |
| PicoSurf | 5 | 300 | 146047 (97.2%) | 142025 (99.4%) | 141185 (99.8%) | 140894 |
| PicoSurf | 5 | 300 | 31351 (98.7%) | 30935 (99.9%) | 30907 (100.0%) | 30891 |
| PicoSurf | 5 | 300 | 109901 (98.3%) | 107999 (99.5%) | 107418 (99.7%) | 107063 |
| PicoSurf | 5 | 300 | 177926 (95.6%) | 170061 (99.8%) | 169738 (100.0%) | 169679 |
| PicoSurf | 5 | 300 | 190749 (97.8%) | 186602 (99.6%) | 185954 (99.8%) | 185511 |

**Supplementary Table 11.** ***Monocore double emulsion picoreactor yield from FACS – FACS tube***

| Surfactant | Concentration (%) | Sort time (sec) | Gate 1 (% passing) | Gate 2 (% passing) | Gate 3 (% passing) | Final |
|---|---|---|---|---|---|---|
| PicoSurf | 5 | 300 | 28397 (98.0%) | 278334 (99.9%) | 27806 (100.0%) | 27798 |
| PicoSurf | 5 | 300 | 30723 (99.3%) | 30505 (100.0%) | 30495 (100.0%) | 30488 |
| PicoSurf | 5 | 300 | 131877 (99.0%) | 130602 (99.8%) | 130405 (100.0%) | 130340 |
| PicoSurf | 5 | 300 | 29113 (98.0%) | 28547 (100.0%) | 28534 (100.0%) | 28533 |

**Supplementary Table 13.** ***Monocore double emulsion picoreactor yield from FACS – FACS tube + Corkscrew***

| Surfactant | Concentration (%) | Sort time (sec) | Gate 1 (% passing) | Gate 2 (% passing) | Gate 3 (% passing) | Final |
|---|---|---|---|---|---|---|
| PicoSurf | 5 | 600 | 299415 (98.0%) | 293347 (99.9%) | 293124 (96.6%) | 283066 |
| PicoSurf | 5 | 600 | 287900 (97.8%) | 281472 (99.9%) | 281231 (97.0%) | 272709 |
| PicoSurf | 5 | 600 | 308296 (97%) | 299024 (99.8%) | 298535 (97.7%) | 291723 |
| PicoSurf | 5 | 600 | 381218 (88.2%) | 336.292 (99.7%) | 335263 (95.6%) | 320653 |

### Supplementary Figures and Legends

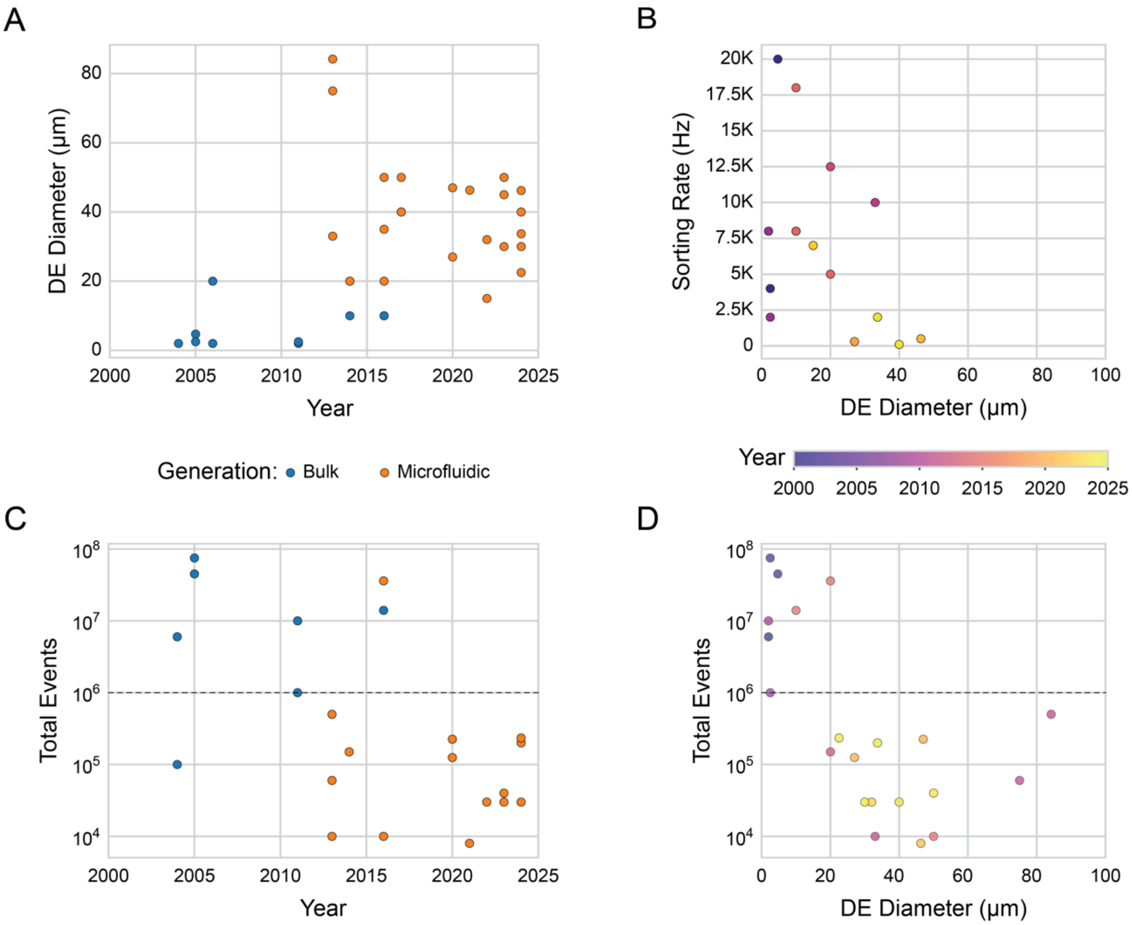


**Figure S1:** ***Analysis of published FACS sortable double emulsion generation methods and throughput.*** Published data for double emulsion (DE) FACS sorting extracted from Supplementary Table 1. **A**) DE diameter *vs.* year of publication. Marker color indicates DE generation method: bulk homogenization or extrusion (blue) or microfluidic droplet generators (orange). **B**) Reported sorting rate *vs.* DE diameter. Marker color indicates publication year (as detailed in colorbar). **C**) $Log_{10}$-transformed total FACS events (e.g. DEs) reported over all publication figures *vs.* year of publication. Marker colors indicate generation method as in A); dashed line indicates 1 million events. **D**) $Log_{10}$-transformed total FACS events *vs.* DE diameter. Marker color indicates publication year (as detailed in colorbar); dashed line indicates 1 million events.

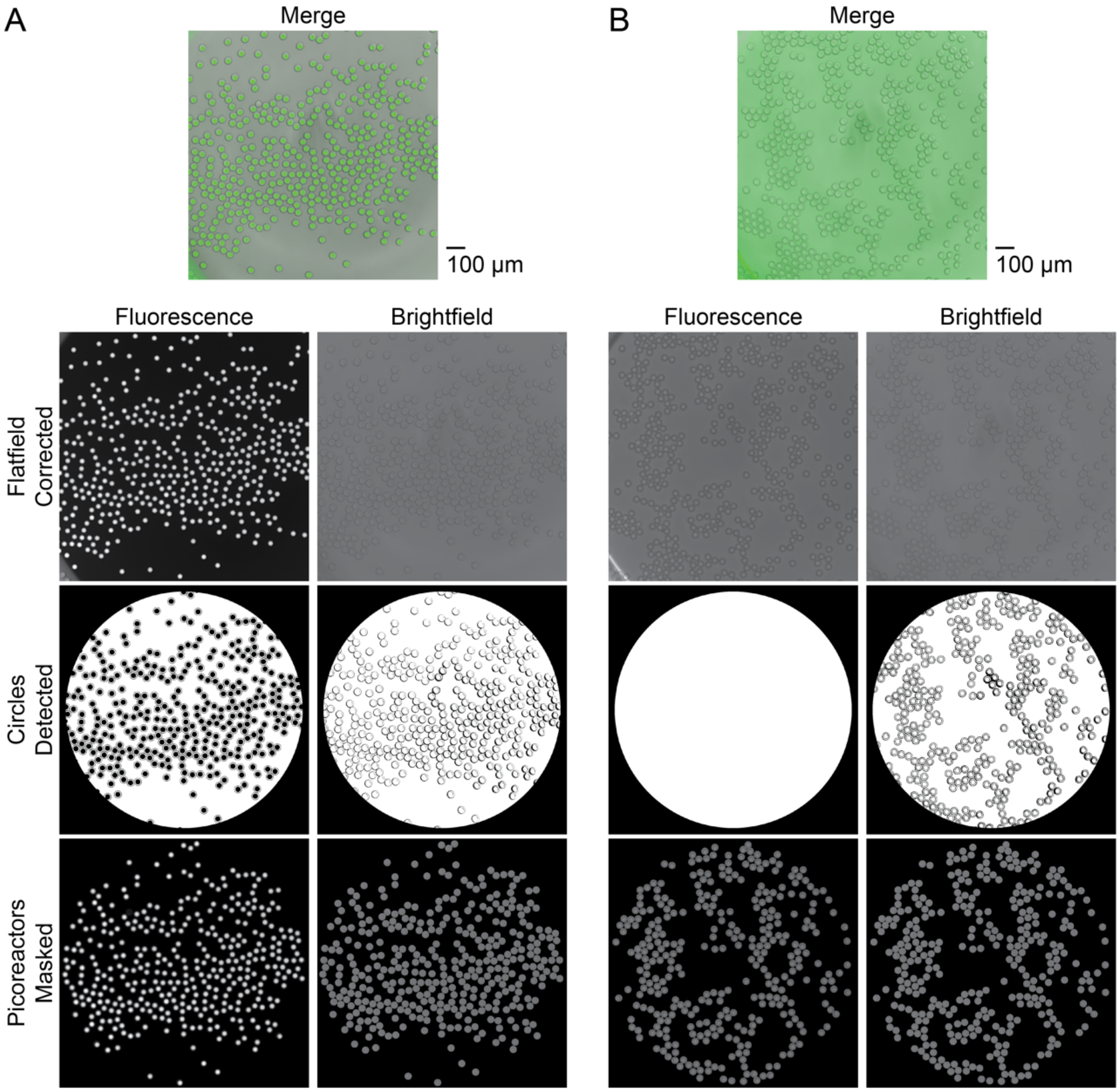


**Figure S2:** ***Automated detection of double emulsions.***
DEs are detected in brightfield and fluorescence images with a custom Python script using the PIL and cv2/OpenCV libraries. Paired fluorescence and brightfield images are clipped and circles corresponding to unique picoreators are detected for masking. DEs can be identified primarily from fluorescence images (**A**) or brightfield images (**B**), dependent on the labeling of the DE. The unique set of DEs detected from both brightfield and fluorescence images are masked from flat field-corrected images and used for analysis. Scale bar: 100 μm.

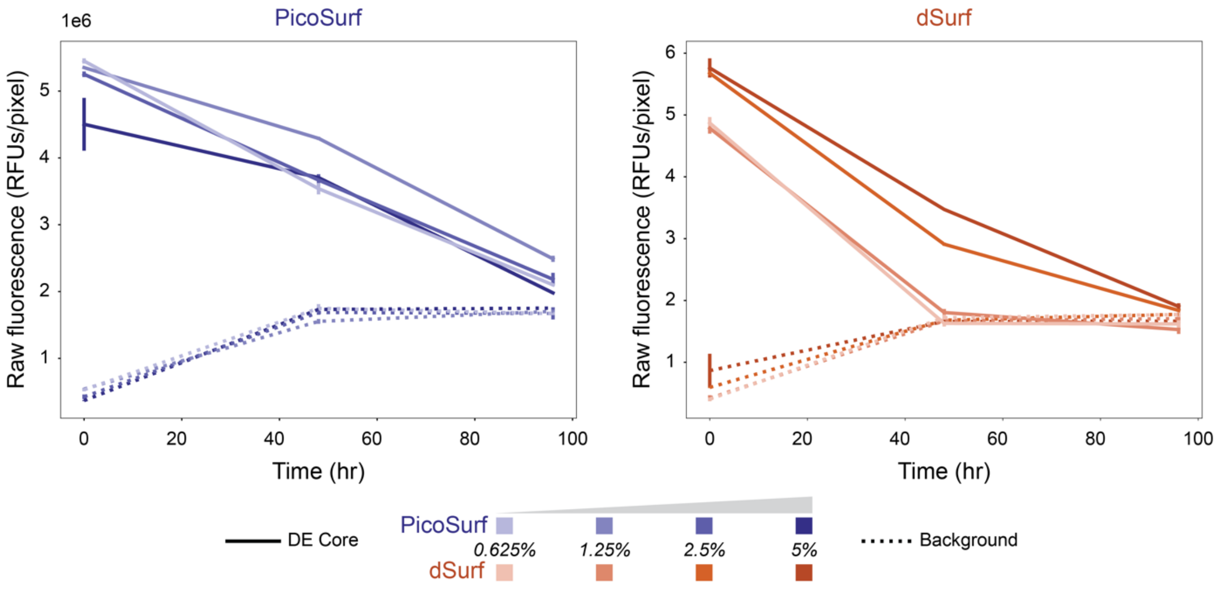


**Figure S3:** ***Raw fluorescence and background intensities for fluorescein-containing double emulsion picoreactors.***
Raw fluorescence intensities for DE cores (solid lines) and background (dashed lines) over all surfactant conditions. PicoSurf (blue) and dSurf (orange) conditions are shaded by the concentration of surfactant as in Figure 1 (see legend, bottom). Per-pixel mean DE core intensities are calculated over a circle covering the center of each automatically detected DE; per-pixel mean background intensities are calculated for an annulus around each core (see **Extended Methods**). Error bars represent standard deviations over three images for all DEs in each image.

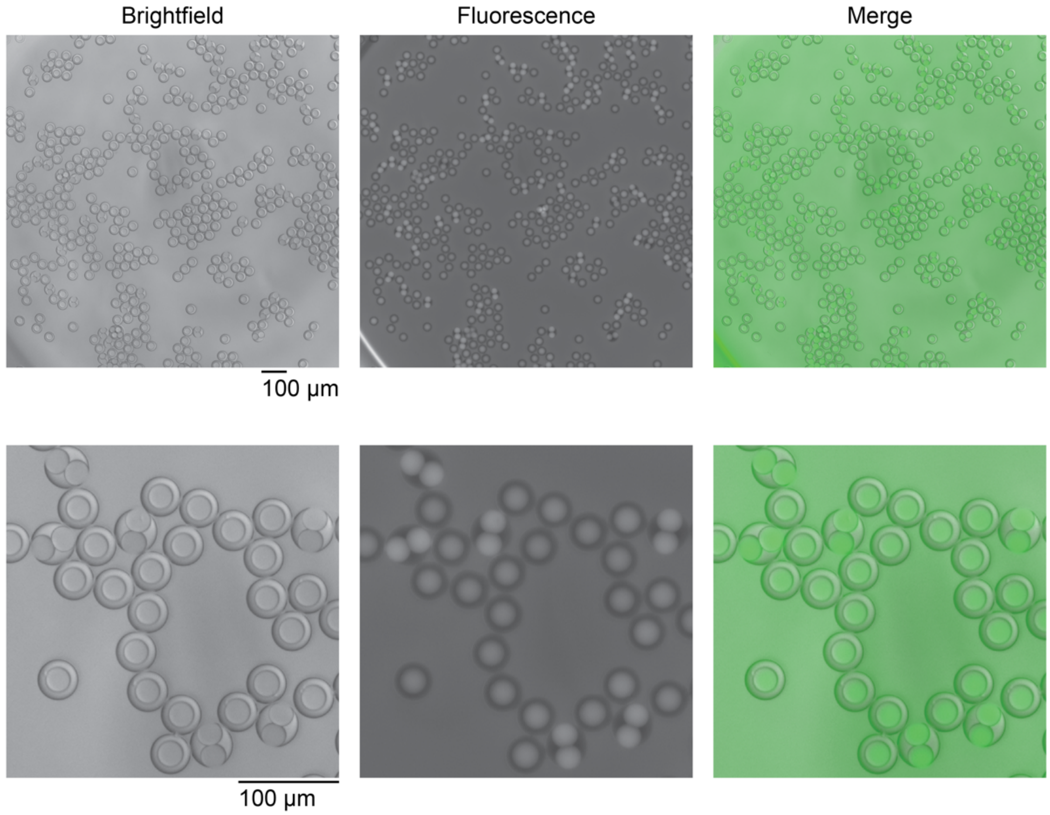


**Figure S4:** ***Merged multicore picoreactors show higher retention of fluorophore in aqueous cores.*** Top: representative brightfield, fluorescence, and merged images for DEs with 0.625% PicoSurf in the oil shell and 20 µM fluorescein in the core after 96 hrs incubation at room temperature. Out of all tested conditions, 0.625% PicoSurf resulted in the highest percentage of merged DEs (see Figure 1C). Scale bar: 100 µm. Bottom: corresponding zoom in for each image. Scale bar: 100 µm.

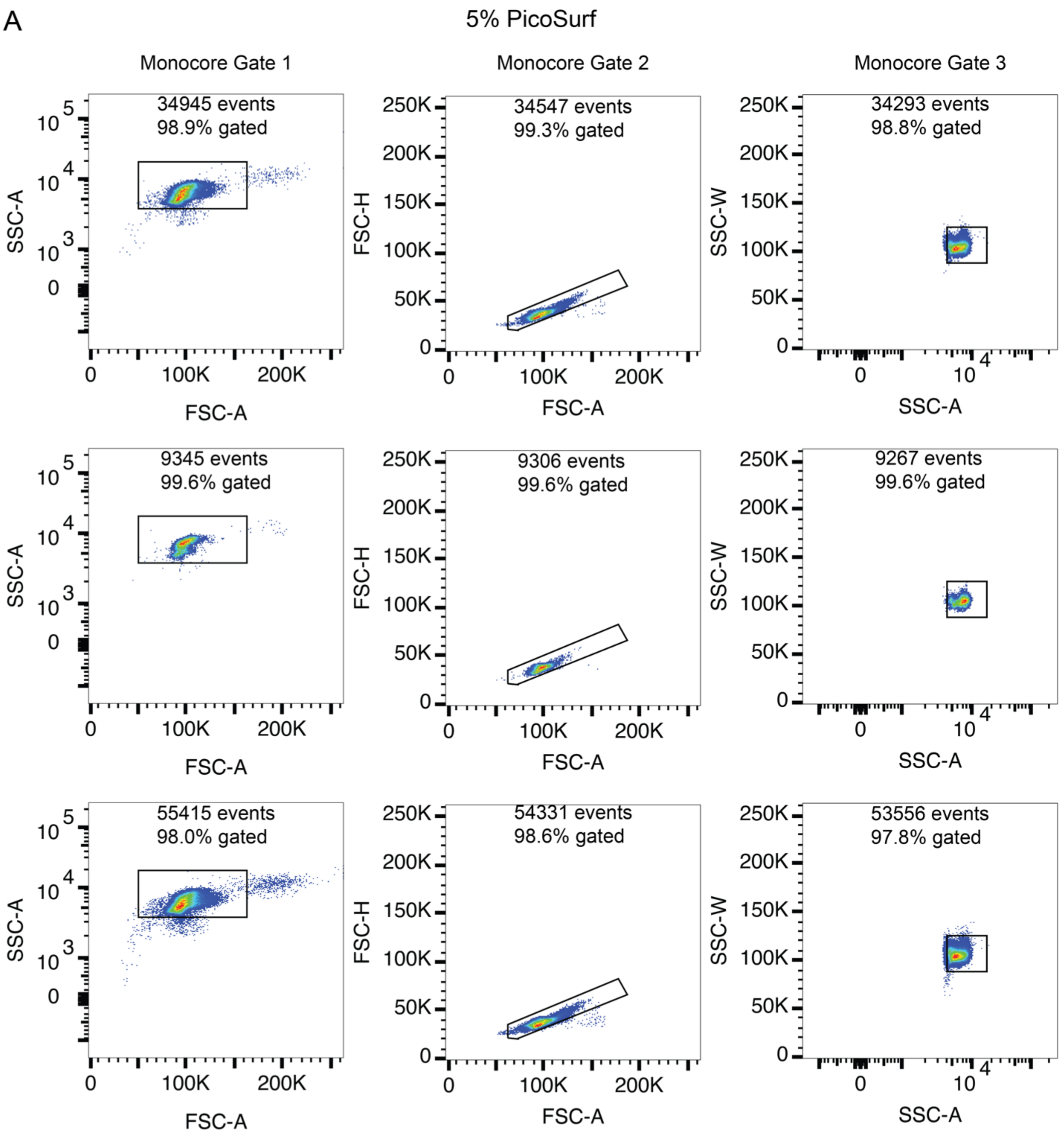


**Figure S5:** ***FACS analysis of double emulsion picoreactor loading and uniformity with PicoSurf surfactant.***
Legend on following page

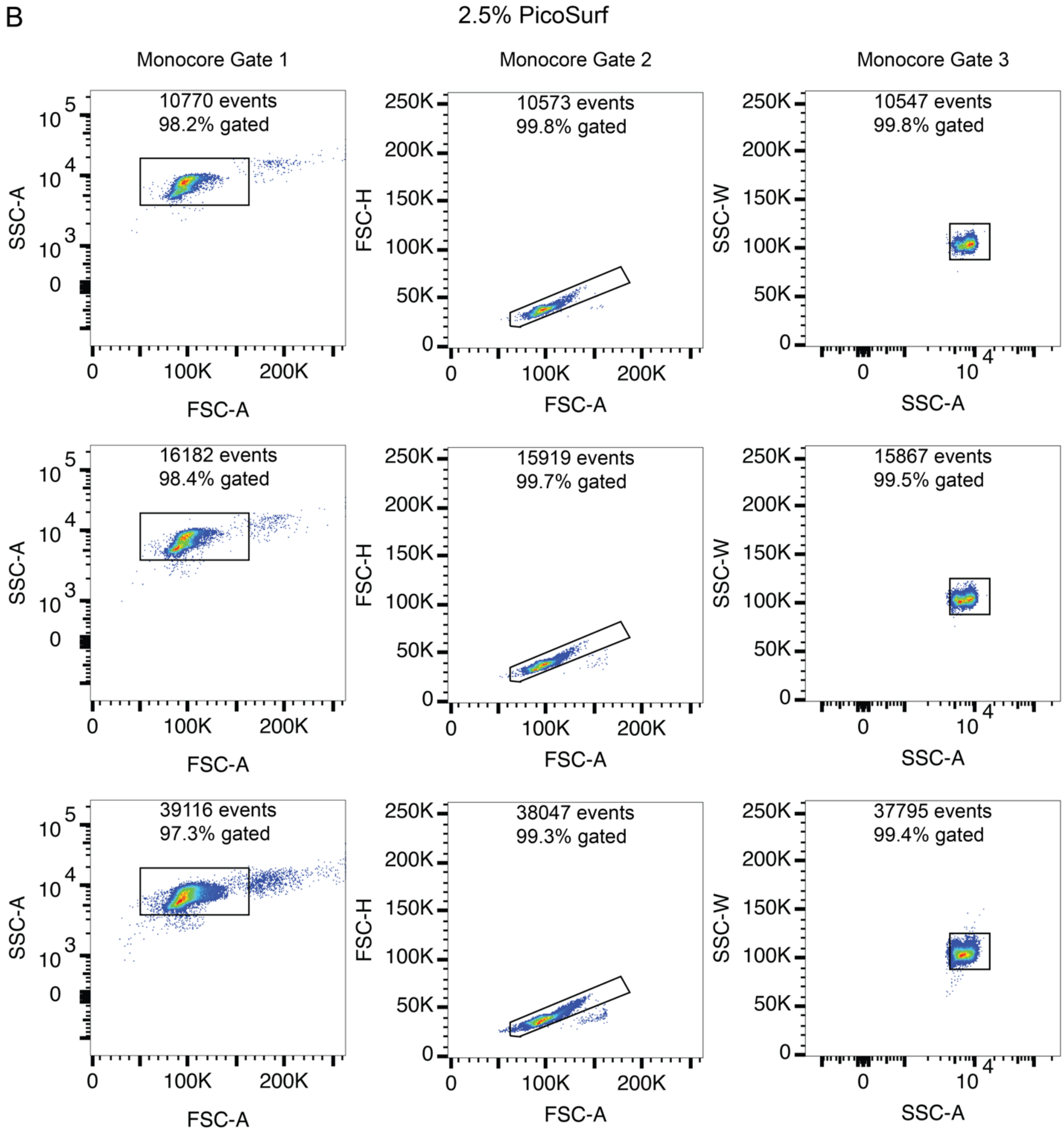


**Figure S5:** ***FACS analysis of double emulsion picoreactor loading and uniformity with PicoSurf surfactant.***
Legend on following page.

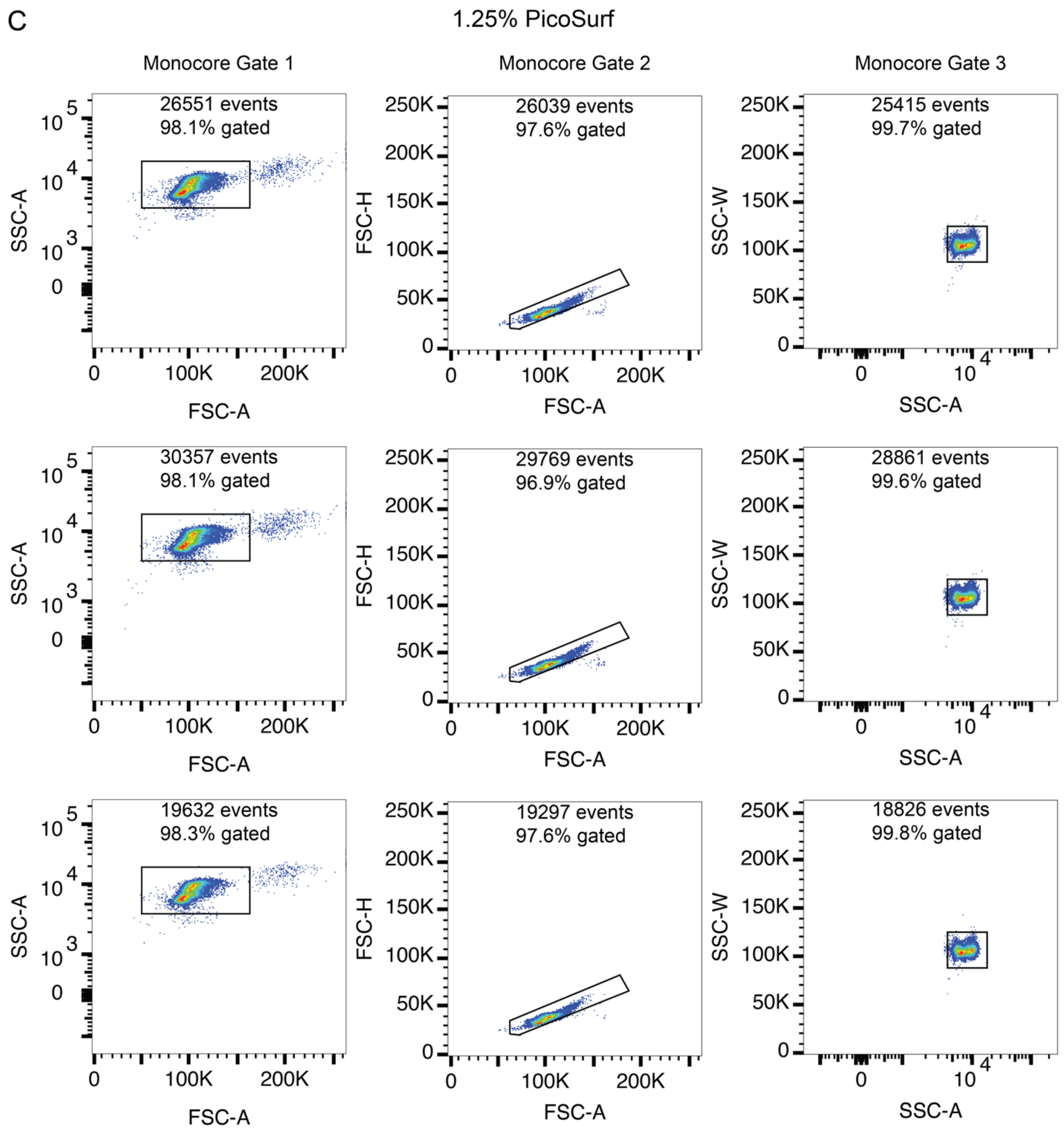


**Figure S5:** ***FACS analysis of double emulsion picoreactor loading and uniformity with PicoSurf surfactant.***

Side-scatter (SSC) and forward-scatter (FSC) plots from FACS analysis of DEs with 5% (**A**), 2.5% (**B**), or 1.25% (**C**) PicoSurf used for analysis in Figure 2. Three replicates are shown for each condition. Monocore DEs are identified through three progressive gates (left-to-right in order of application): forward-scatter area (FSC-A) vs. side-scatter area (SSC-A), FSC-A vs. forward-scatter height (FSC-H), and SSC-A vs. side-scatter width (FSC-W).

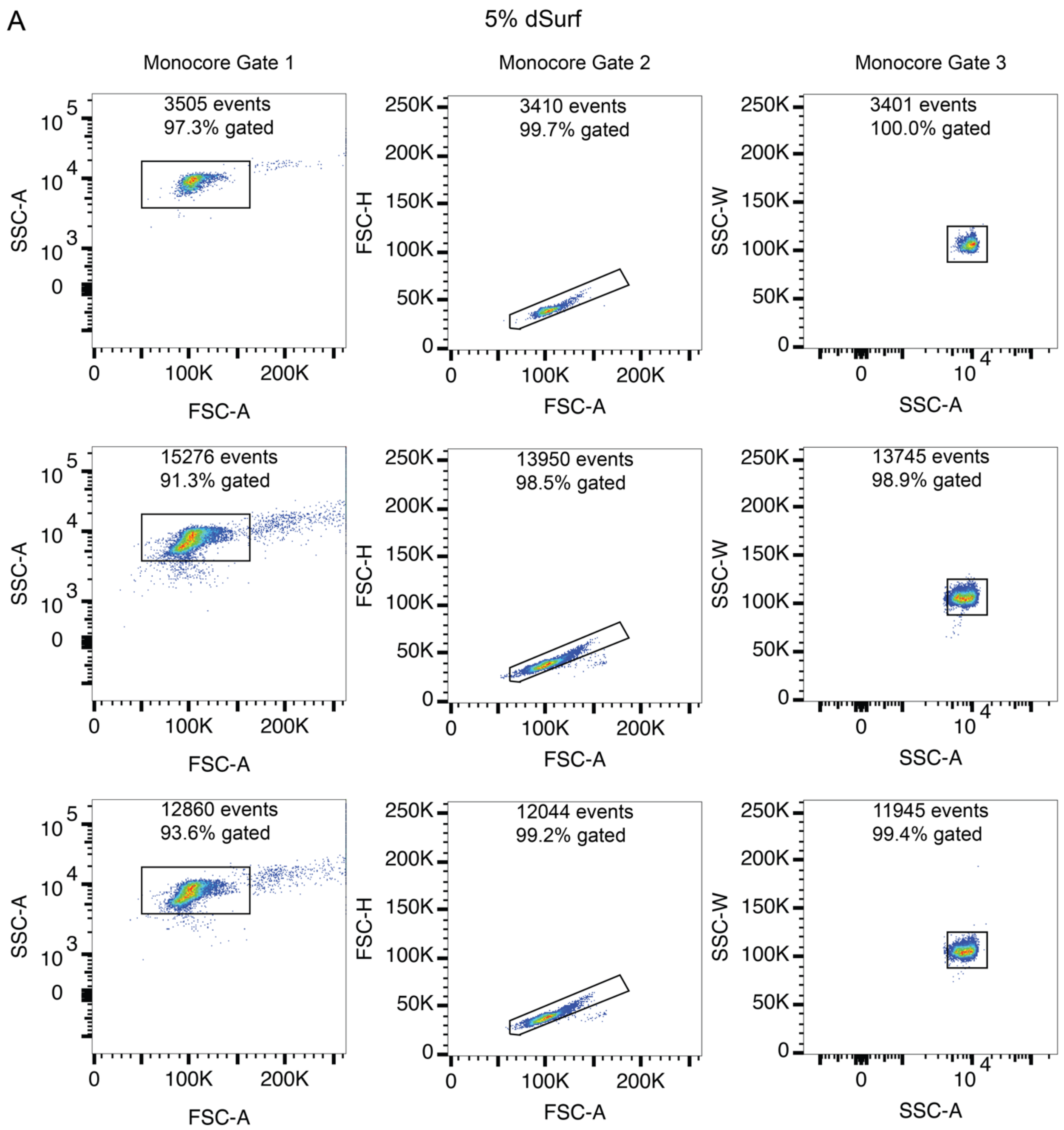


**Figure S6:** ***FACS analysis of double emulsion picoreactor loading and uniformity with dSurf surfactant.***

Legend on following page

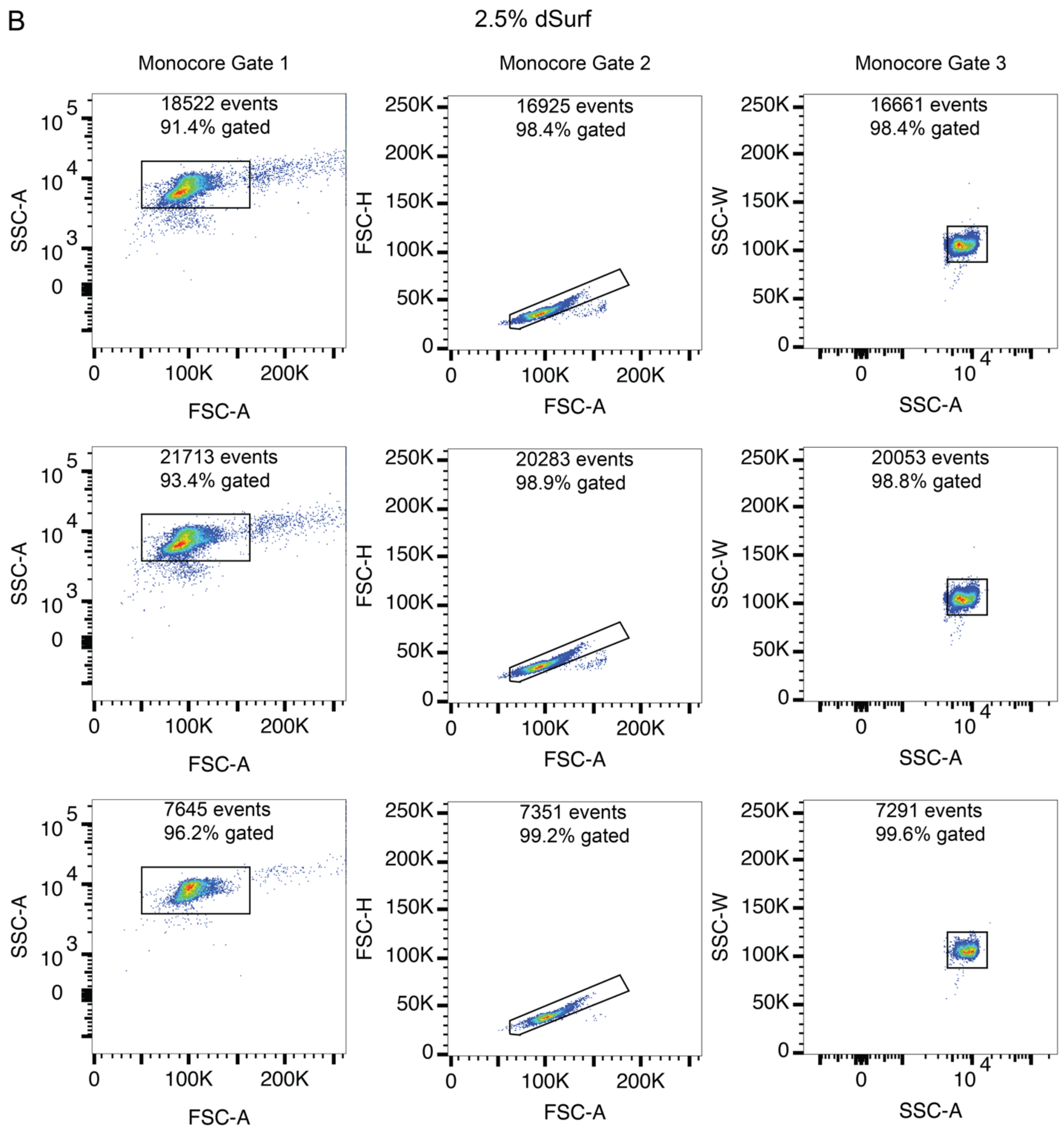


**Figure S6:** ***FACS analysis of double emulsion picoreactor loading and uniformity with dSurf surfactant.***

Legend on following page.

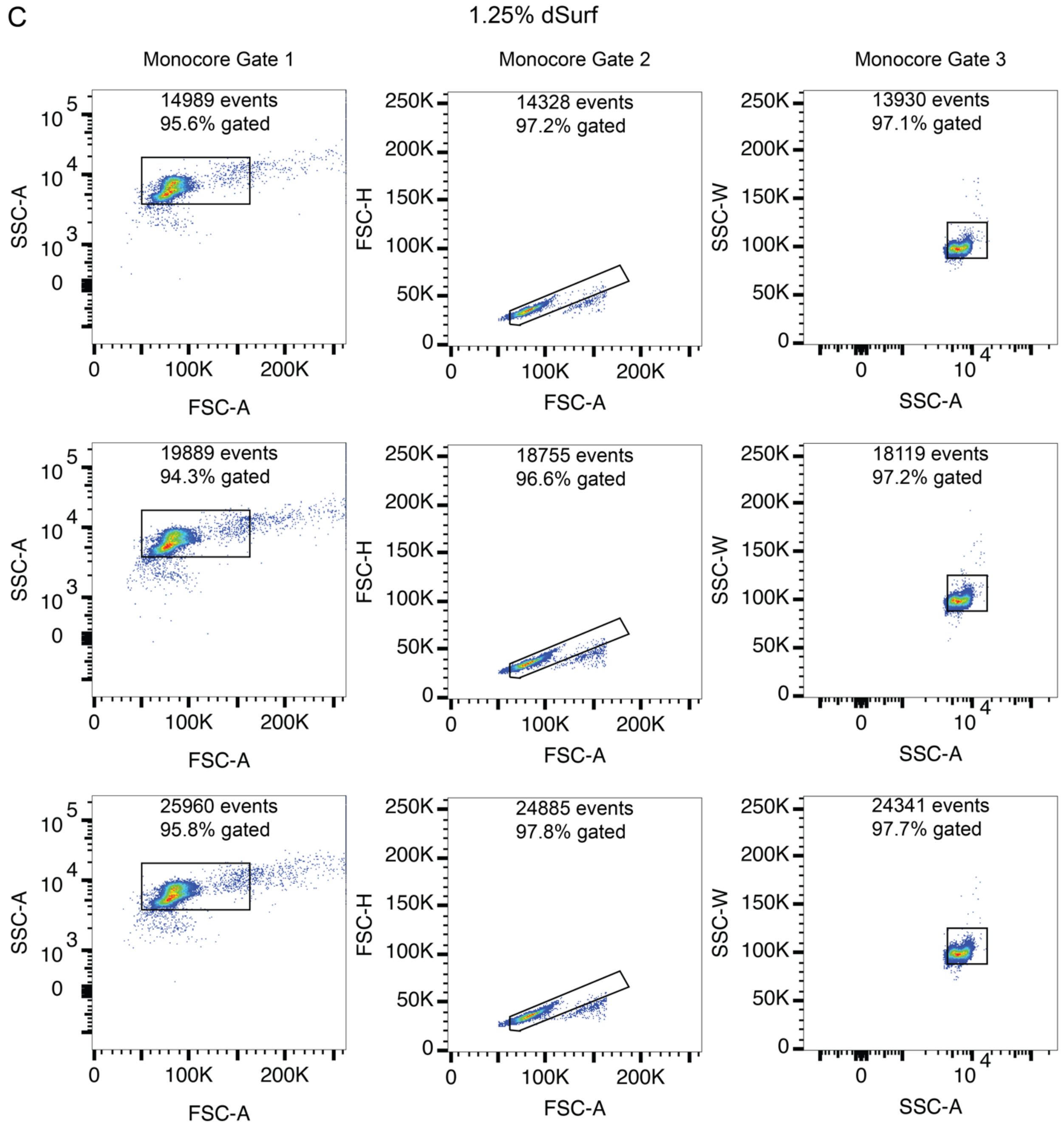


**Figure S6:** ***FACS analysis of double emulsion picoreactor loading and uniformity with dSurf surfactant.***
SSC and FSC plots from FACS analysis of DEs with 5% (**A**), 2.5% (**B**), or 1.25% (**C**) dSurf used for analysis in Figure 2. Three replicates are shown for each condition. Monocore DEs are identified through three progressive gates (left-to-right in order of application): FSC-A vs. SSC-A, FSC-A vs. FSC-H, and SSC-A vs. FSC-W, as described in Figure S3.

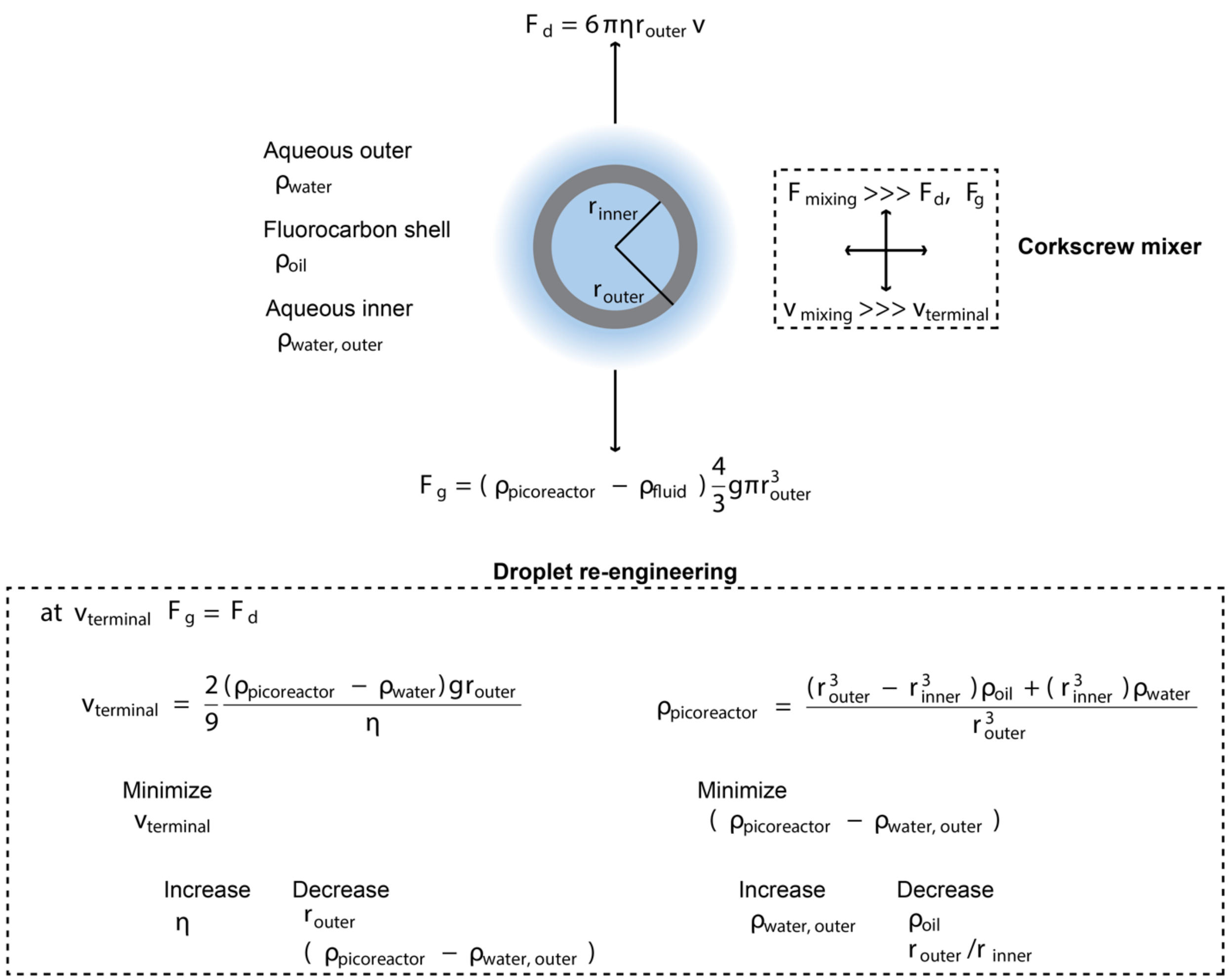


**Figure S7:** ***Stokes law model of double emulsion picoreactor settling.***
A DE is modeled as a hard sphere falling in a solution of pure water. To simplify this conceptual analysis, the impact of buffers and salts on water density and the deformability of the DE are not considered. DE settling terminal velocity is achieved when the forces of gravity and drag cancel each other out. Gravitational force ($F_g$) is a function of difference in mass for the DE relative to the mass of outer fluid for an equal size. DE mass is a function of the size of the DE ($\frac{4}{3}\pi r_{outer}^3$), the thickness of the oil shell ($r_{outer} - r_{inner}$), and the density of the oil ($\rho_{oil}$). The force of drag ($F_d$) for a falling hard sphere is dependent on the size of the DE ($r_{outer}$), the viscosity of the outer fluid ($\eta$), and the velocity (v) of the falling particle. $v = v_{terminal}$ when $F_g = F_d$. A stable suspension of DEs would have a v =0, so to minimize v, multiple parameters can be decreased ($r_{outer}$ or $\rho_{picoreactor} - \rho_{water,outer}$) or increased ($\eta$). Also, when $F_{mixing} >> F_g$ and $F_d$, then $V_{mixing} >> V_{terminal}$ and the DEs will remain resuspended.

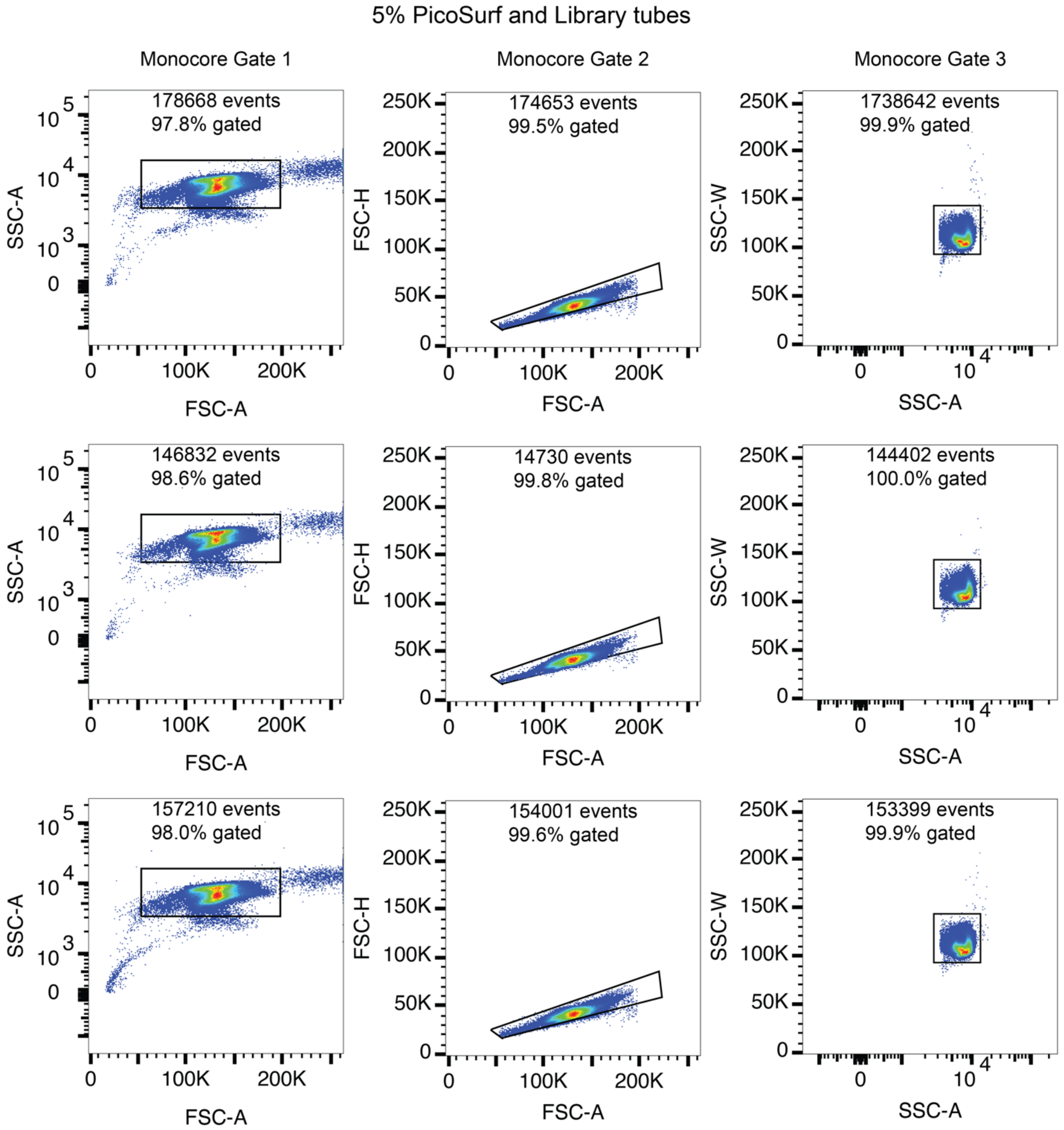


**Figure S8:** ***FACS analysis of double emulsion picoreactor loading and uniformity with library tube loading.***

Legend on following page.

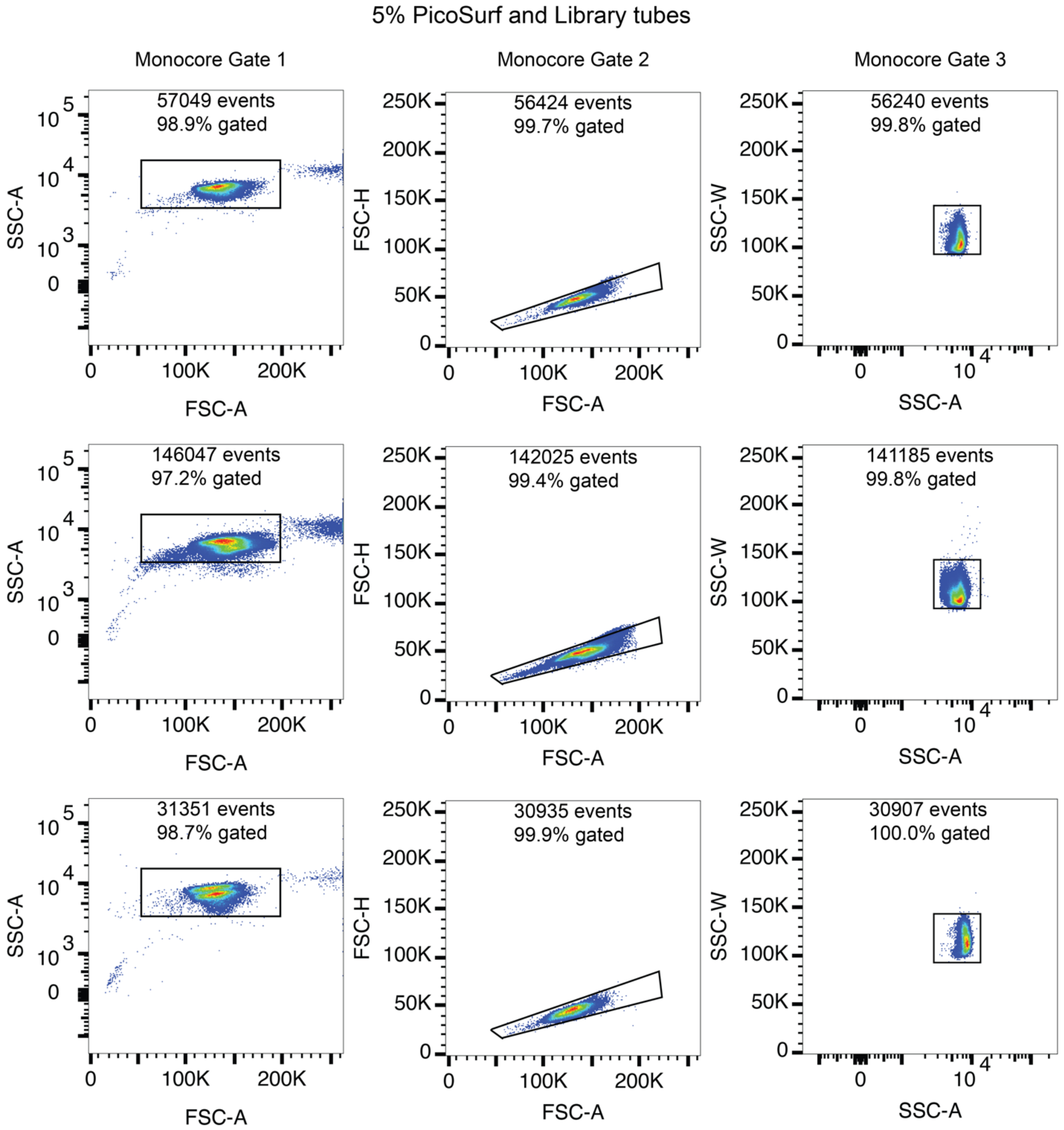


**Figure S8:** ***FACS analysis of double emulsion picoreactor loading and uniformity with library tube loading.***

Legend on following page.

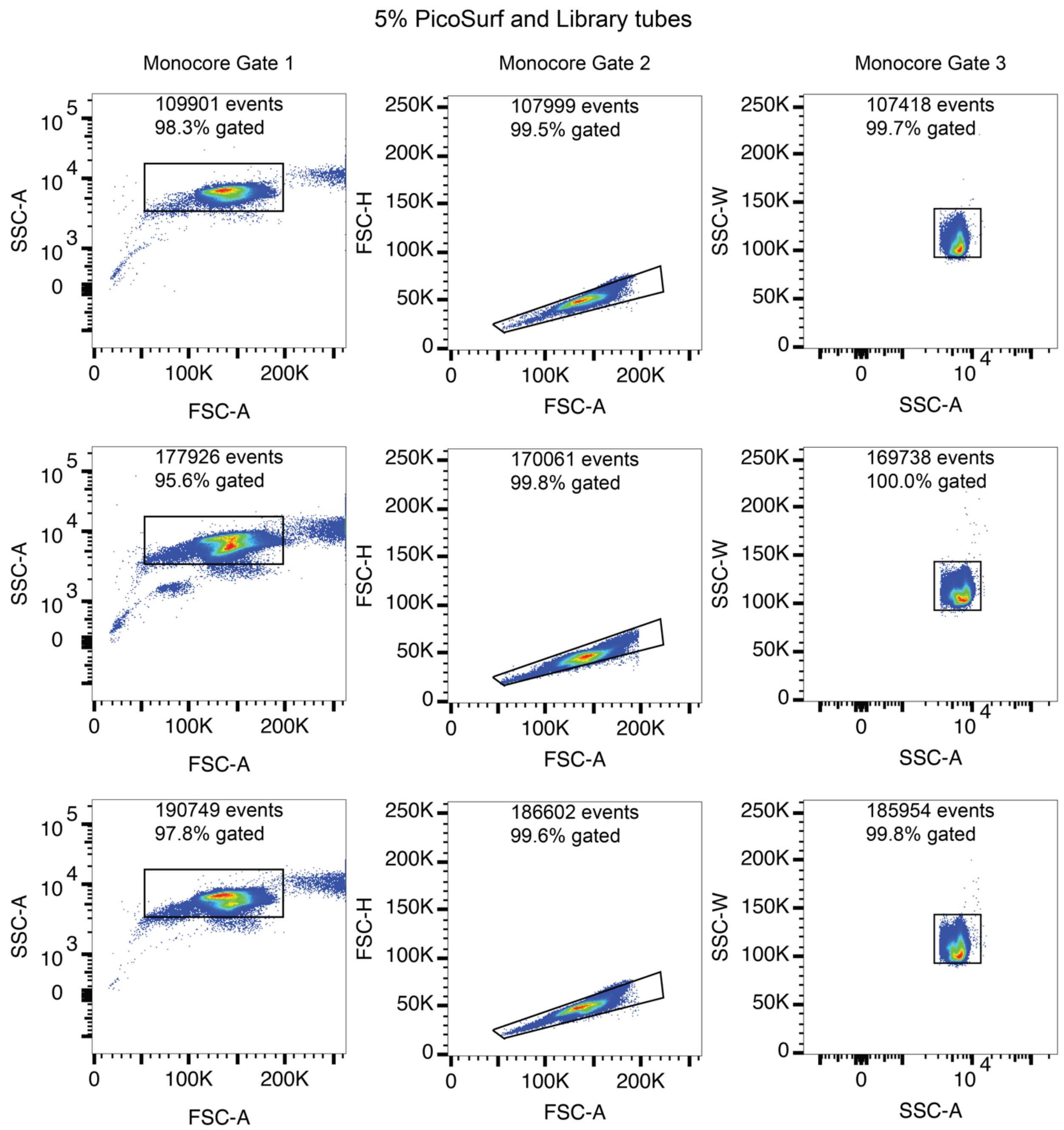


**Figure S8:** ***FACS analysis of double emulsion picoreactor loading and uniformity with library tube loading.***

SSC and FSC plots from FACS analysis of DEs with 5% PicoSurf loaded with library tubes and used for analysis in Figure 3. Nine replicates are shown for each condition. Monocore DEs are identified through three progressive gates (left-to-right in order of application): FSC-A vs. SSC-A, FSC-A vs. FSC-H, and SSC-A vs. FSC-W, as described in Figure S3.

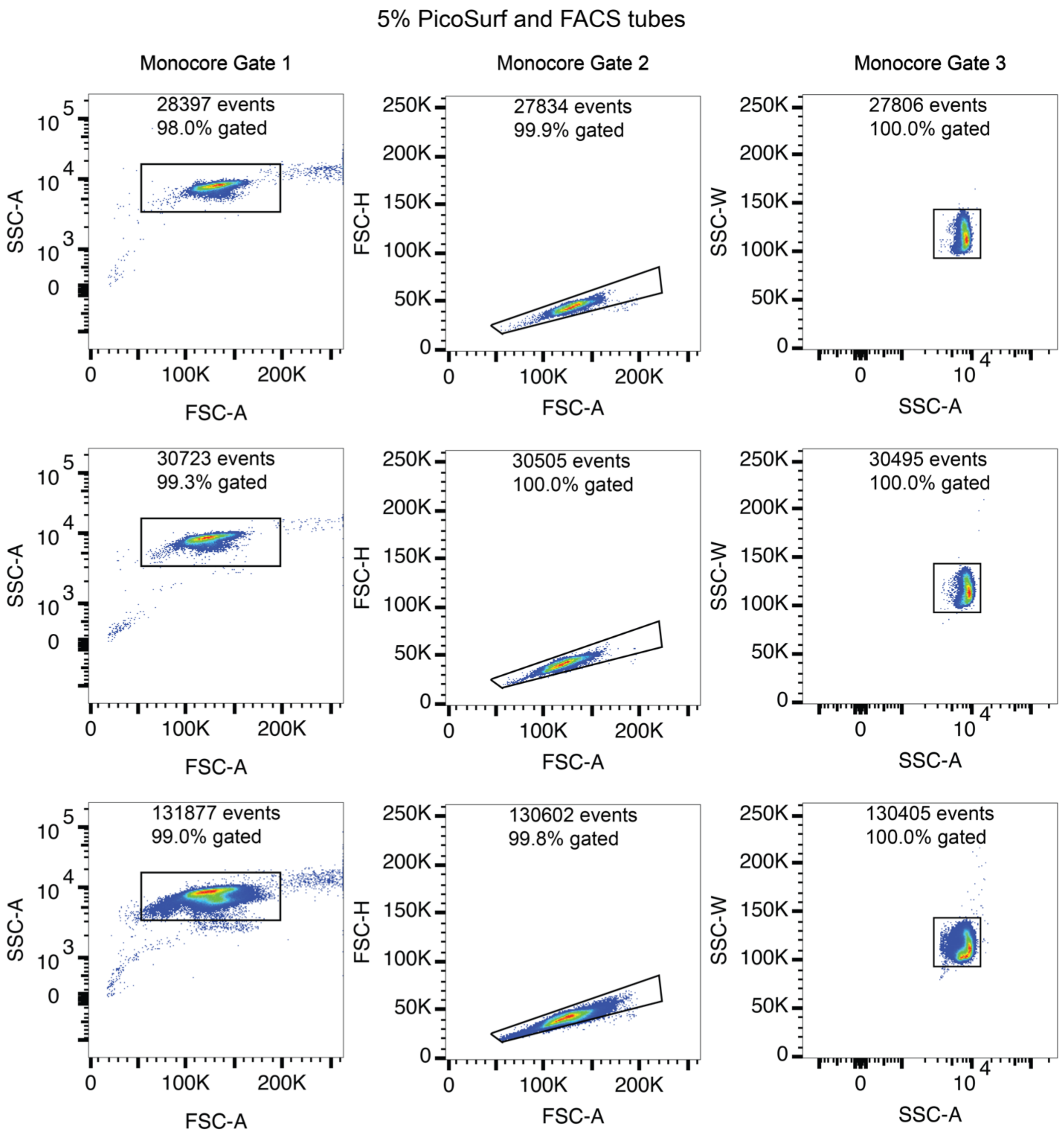


**Figure S9:** ***FACS analysis of double emulsion picoreactor loading and uniformity with FACS tube loading.***
Legend on following page.

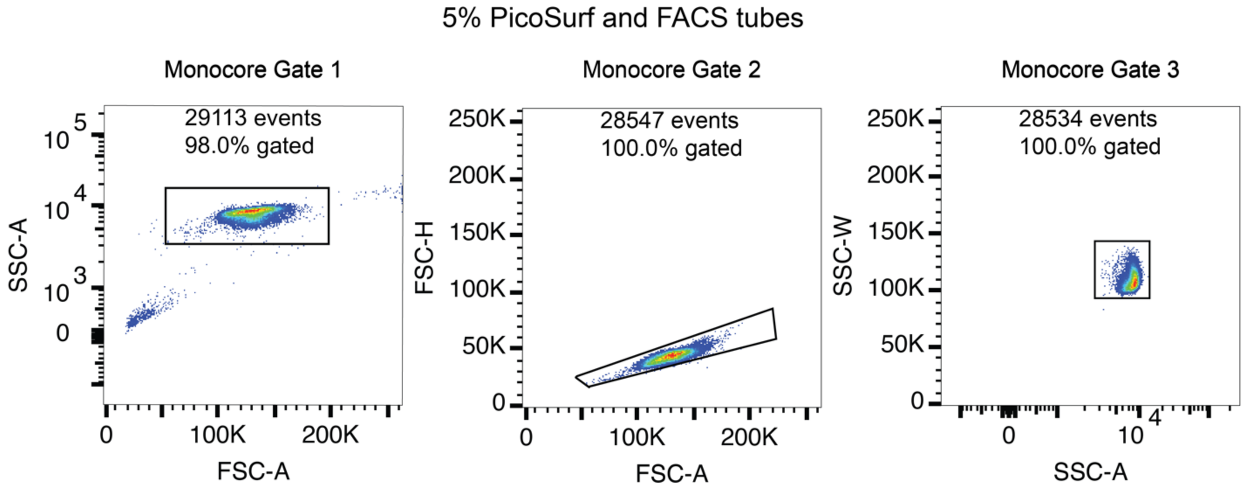


**Figure S9:** ***FACS analysis of double emulsion picoreactor loading and uniformity with FACS tube loading.***
SSC and FSC plots from FACS analysis of DEs with 5% PicoSurf loaded with FACS tubes and used for analysis in Figure 3. Four replicates are shown for each condition. Monocore DEs are identified through three progressive gates (left-to-right in order of application): FSC-A vs. SSC-A, FSC-A vs. FSC-H, and SSC-A vs. FSC-W, as described in Figure S3.

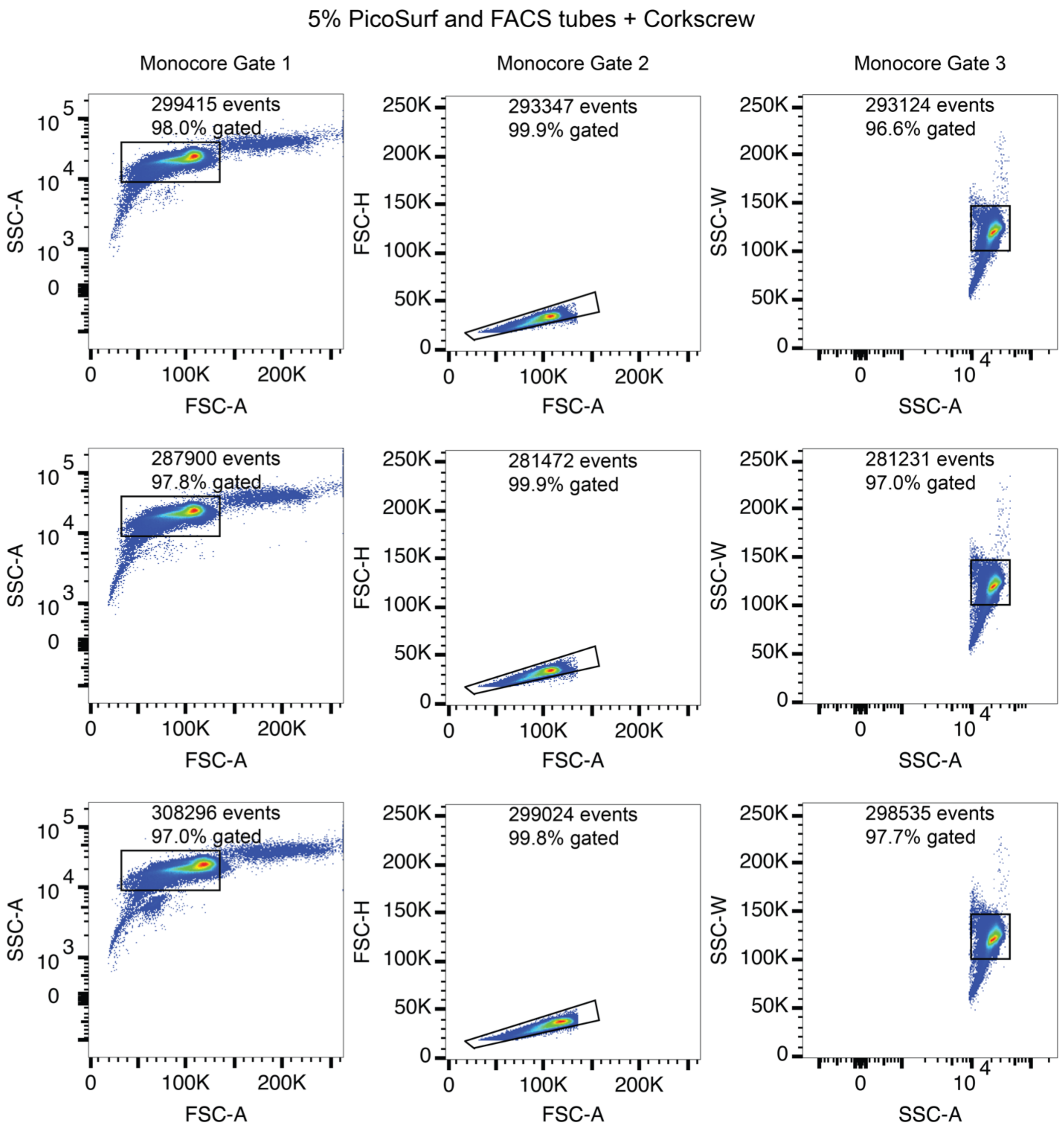


**Figure S10:** ***FACS analysis of double emulsion picoreactor loading and uniformity with corkscrew mixer loading.***

Legend on following page.

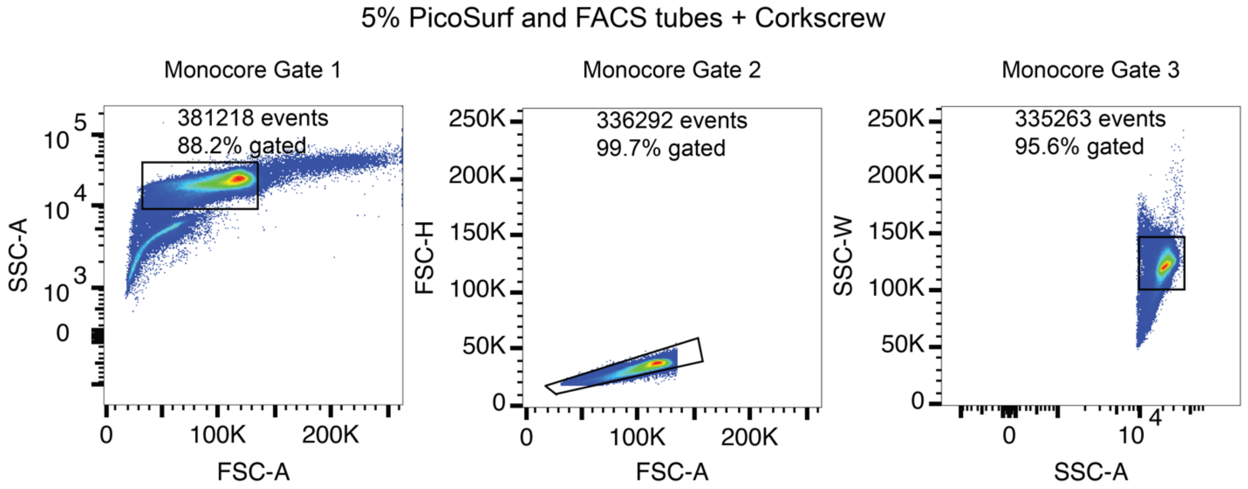


**Figure S10:** ***FACS analysis of double emulsion picoreactor loading and uniformity with corkscrew mixer loading.***

SSC and FSC plots from FACS analysis of DE with 5% PicoSurf loaded with FACS tubes using the corkscrew mixer and used for analysis in Figure 3. Four replicates are shown for each condition. Monocore DE are identified through three progressive gates (left-to-right in order of application): FSC-A vs. SSC-A, FSC-A vs. FSC-H, and SSC-A vs. FSC-W, as described in Figure S3.

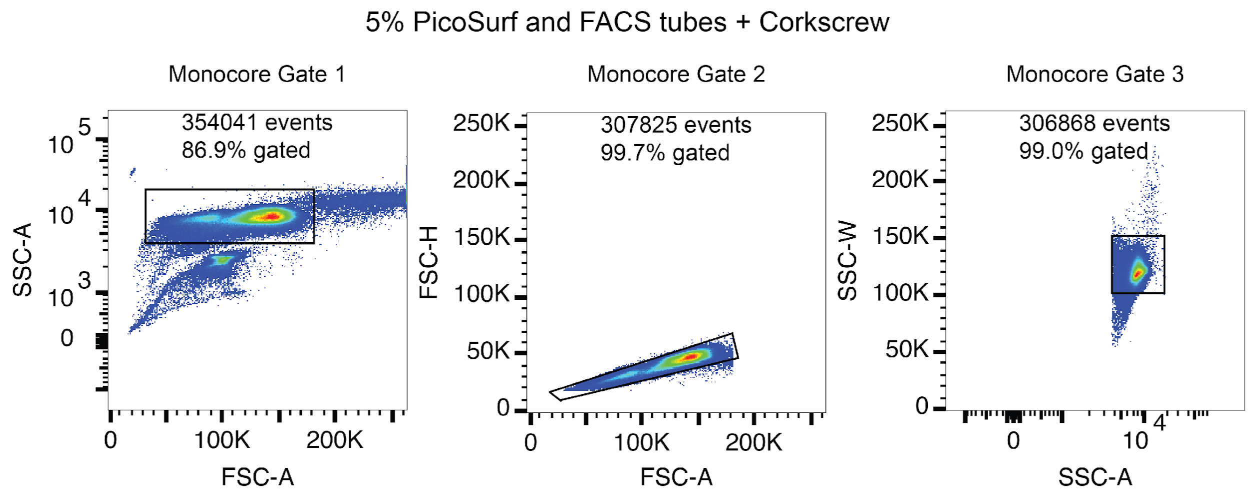


**Figure S11:** ***FACS analysis of double emulsion picoreactor loading and uniformity during rare population selection with corkscrew mixer loading.***
SSC and FSC plots from FACS analysis and sorting of DEs with 5% PicoSurf loaded with FACS tubes using the corkscrew mixer and used for analysis in Figure 4. Four replicates are shown for each condition. Monocore DEs are identified through three progressive gates (left-to-right in order of application): FSC-A vs. SSC-A, FSC-A vs. FSC-H, and SSC-A vs. FSC-W, as described in Figure S3.

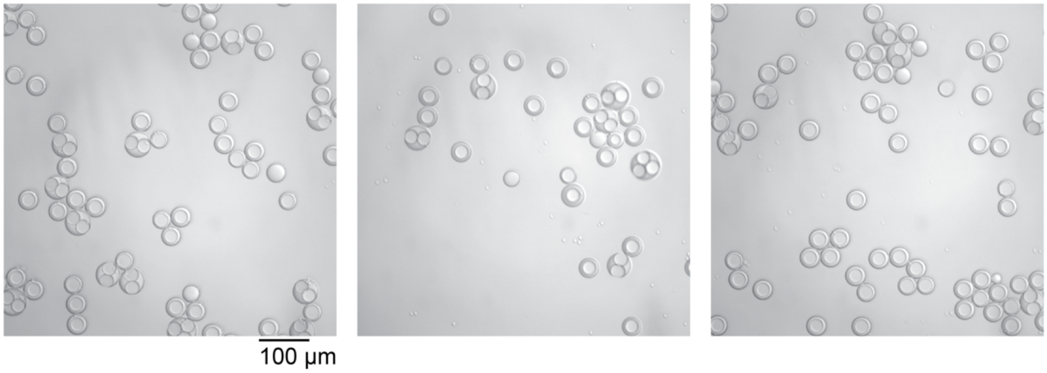


**Figure S12:** ***Sample agitation with corkscrew mixer results in merged double emulsion picoreactors.*** Three representative images of DE samples remaining in the FACS tube after the sorting analysis in Figure 4. Scale bar: 100 μm.

### Extended Methods

#### *Operating microfluidic devices*

Preparing stock reagents

5% PicoSurf in HFE7500 (SphereFluidics, cat. # C022) was used as provided. Three serial 2-fold dilutions were prepared by mixing 2 mL PicoSurf in HFE7500 with 2 mL of HFE7500 (Novec7500, 3M, cat. # 7100025016) in a 20 mL vial (Wheaton, cat. # 986741). After each dilution, the vial was vortexed for 3 minutes at 3000 rpm. 5% dSurf was prepared by measuring 0.5 g of Honey dSurf (Fluigent, cat. # OdSurf_5G) in a 20 mL vial, adding 10 mL of HFE7500, and vortexing for 3 minutes at 3000 rpm. Three serial 2-fold dilutions were prepared by mixing 2 mL dSurf in HFE7500 with 2 mL of HFE7500. After each dilution, the vial was vortexed for 3 minutes at 3000 rpm. All droplet oil solutions were filtered with a syringe-driven 0.22 µm filter (Millipore Sigma, cat. # SLGV033RB). 20 mM fluorescein was prepared by dissolving 6.6 mg fluorescein powder (Fluka, cat. # 46955) in 1 mL of DMSO (Sigma Aldrich, cat. # D8418).

PDMS device generation

Droplet generators were used as designed in Brower et al., 2020[9]. CAD files for photolithography exposure masks are available from the Open Science Framework (OSF) repository for that reference: https://osf.io/h4sr9/. Briefly, molds for device features were generated on silicon wafers using photolithography. PDMS (MG Chemicals, cat. # RTV615) was cast onto a feature mold and a blank silicon water; after a soft bake (80˚C, 12-15 min), the PDMS was released from the mold. Holes were punched for inlet and outlet ports with a 1 mm biopsy punch (Robbins Instruments, cat. #RBP-10), the PDMS pieces were cleaned with Scotch tape and the featured and blank PDMS slabs were joined and baked at 80˚C for 48-56 hrs. AutoCAD files and photolithography protocols for masks used in this study are available in the OSF repository for this manuscript at https://osf.io/jzyve/.

Generating double emulsion picoreactors

FACS sortable double emulsions were generated as previously described using a set-up that included the PDMS devices, 4 syringe pumps (Harvard Apparatus, cat. # 70-4511), and a zoom power optical microscope. Aqueous solutions were prepared and filtered with a syringe-driven 0.22 µm filter. Droplet oil was used for the fluorinated oil shell as generated above. For the inner solution (PBS pH 7.4 (Gibco, cat. # 10010031) + 1% Tween 20, both inlets 1 on diagram), 1 g of Tween 20 (Fisher BioReagents, cat. # BP337) was measured in a graduated cylinder. For the outer solution (PBS pH 7.4 + 1% Tween 20 + 2% Pluronic F68, inlet 3 on diagram), 1 g of Tween 20 and 2 g of Pluronic F68/Kolliphor P 188 (Millipore Sigma, cat. # K4894) were measured in a graduated cylinder. For each, the volume was brought up to 100 mL with PBS pH 7.4. A stir bar was added to each graduated cylinder, and the opening was sealed with Parafilm (Bemis, cat. # PM-996). The surfactants were dissolved by overturning and stirring, then the solutions were sterile-filtered into fresh PETG storage bottles (Nalgene, cat. # 342020-0030). For labeled inner solution with 20 µM fluorescein, 10 mL of PBS pH 7.4 + 1% Tween 20.

For DE generation, input solutions were loaded into disposable Luer lock syringes: inner solution in two 1 mL syringes, dSurf oil in a 3 mL or 5 m syringe, and outer solution in a 10 mL or 30 mL syringe (Becton Dickinson, cat. # 309628, 309657, 309646, 302995, 302832). Bubbles were removed from each solution by tapping and ejection, then each syringe was capped with a 27-gauge stainless steel dispensing needle (McMaster-Carr, cat. # 75165A688). The needles were inserted into the open end of polyethylene medical tubing (Scientific Commodities Inc., cat. # BB31695-PE/2). The syringes were installed into the pumps, and the tubing was cut sufficiently long to reach the PDMS device on the stage. The device was centered in the microscope field of view, and the free ends of the tubing were inserted directly into the ports of the PDMS device after trimming the tubing to reduce slack. The tubing was removed, and the right-hand side of the device was plasma treated to render it hydrophilic. The ports for inner and oil phases were covered with Scotch tape, and the PDMS device array was placed in a plasma cleaner (Harrick Plasma, cat. # PDC-001) with a dry scroll pump (Agilent cat. # IDP3B01), a Type 0536 TC vacuum gauge (Agilent cat. # L6141303), and a vacuum gauge monitor (Harrick Plasma, cat. # PDC-VCG). The chamber was vacated, and the three-way valve was opened to ambient air with the regulator valve that maintained pressure at approximately 400 mbar. Plasma treatment was performed under "high" setting (30 watts) for 10-12 minutes. After deactivating plasma coils, the chamber was repressurized to ambient pressure and the PDMS device array was removed.

After plasma treatment, 45 µm devices were operated as follows. Fluid solutions were ejected sufficient to fill the tubing with solution and eliminate air bubbles. An outlet line was connected to the device and the free end was inserted into a fresh 2 mL tube (outlet 4 on diagram) The device was re-centered on the microscope, and the outer solution line was connected to the device and the outer solution was flowed through the port for 30 sec at a flow rate of 3000 µL/hr. The oil input was set to a flow rate of 400 µL/hr and the tubing was connected to the port. Once the oil was visible in the second first flow focuser the flow rate was cut back to 200 µL/hr and adjusted to between 125 µL/hr. Finally, the two inner inputs were set to flow rates of 100 µL/hr each and the lines were connected to the device. When the inner solution was visible in the first flow focuser, each flow rate was decreased to 25 µl/hr. The outer flow rate was gradually increased to 3500 µL/hr. Flow rates were periodically adjusted to maintain stable DE generation or to alter the DE geometry.

***Imaging double emulsion picoreactors***

DEs were imaged on an inverted light microscope (Nikon, Eclipse Ti). Samples consisting of double emulsions were resuspended in solution by overturning the sample tube, and 10 µL of undiluted solution was pipetted onto a Countess chamber slide (ThermoFisher Scientific, cat. # C10228). The light microscope was controlled with a distribution of the MicroManager software package. Images were taken with a combination of brightfield (Semrock, BRFLD-A-NTE-ZERO) and green fluorescence (Chroma, 96226, Ex.: AT480/30x, Dichroic mirror: AT505DC, Em.: AT535/40m). Exposure times were 10 ms for bright field and 500 ms for green fluorescence (fluorescein). Illumination was provided by a SOLA SE light engine through a 3mm liquid light guide (Lumencor, cat # SOLA SE 5-LCR-SA). A 10x objective lens was used for all images (Nikon, cat# MRD00100). Images were captured on an sCMOS camera (Andor, Zyla 4.2 Plus, VSC-06278). All images were recorded at 1x1 binning.

DE images were analyzed with a custom Python script using the PIL and OpenCV libraries. Scripts and data are available at https://osf.io/jzyve/. Briefly, corresponding brightfield and fluorescence images are provided as input and flatfield corrected relative to a standard set of reference images for each channel. Brightfield images are subject to a binary threshold filter and vignette masking before individual monocore DEs are identified with the HoughCircle method from the OpenCV library. Binary thresholding and vignette masking parameters were manually modified for each image to ensure accurate detection. Prior to monocore detection, multicore particles were manually detected in ImageJ and output files containing their x,y positions and their diameters were passed to the python script. These regions were masked out from automated detection, and the number of cores in the multicore DEs was approximated from the calculated volume of the aggregate and an expected volume for a monocore DE. For each condition and timepoint, the percentage of monocore DEs was calculated in each of three images and reported as the mean percentage ± standard deviation.

Fluorescence intensity within each DE was quantified within a user-defined radius from the center of the DE and then normalized to a per-pixel average intensity by dividing the integrated fluorescence by the area of the circle. The background intensity was calculated for each individual DE by calculating the per-pixel average intensity in a ring around the center circle over which fluorescence intensity is calculated. The ring has a user-defined thickness and a user-defined offset from the center circle. For each condition and timepoint, fluorescence intensities were calculated for all monocore particles in each of three images. The mean value was calculated for each image, and then the mean value and standard deviation of the mean intensity was calculated over all three images. For each condition and timepoint, the values were reported as the percent fluorescent intensity relative to the three-image mean from the 0 hr timepoint.

#### *FACS sorting microfluidic double emulsion picoreactors*

3D printing a corkscrew vertical mixer

The 3D corkscrew component installed on a FACSAria II (BD) to facilitate DE sorting was custom-designed using AutoCAD software (Autodesk). The direction of rotation varies by instrument, so designs for both right-handed and left-handed corkscrews have been uploaded to Thingiverse and is available for other users under a Creative Commons license (https://www.thingiverse.com/thing:6675740). The corkscrew device was printed using a high-resolution stereolithography 3D printer Anycubic Photon Mono 4K (Anycubic). The slicing of the corkscrew was performed in AnycubicPhotonWorkshop software (Anycubic), with a layer thickness of 50 µm, a normal exposure time of 2 seconds, and a bottom exposure time of 40 seconds. The corkscrew was printed using an Anycubic 3D printing UV sensitive Resin (US, RPTTO2309C0601), while being horizontally oriented.

After printing, the central hole of the corkscrew was cleared using a heated 0.8 mm needle (McMaster, cat M925, Dispos. LL TW Probe Needle, 23x1/2) to ensure an unimpeded tunnel. Finally, the resin was cured by exposure to UV light for 15 minutes using the Anycubic Wash&Cure 2.0 setup.

FACS calibration, analysis, and sorting

DE sorting was conducted using a FACSAria II cell sorter (BD) following an optimized calibration workflow on. Laser delays settings were established with 32 µm AccuCount Ultra Rainbow calibration beads (Spherotech, cat. # ACURFP2.5-300-1). The forward scatter threshold was set to ½ of the of the mean FSC-H, the blue laser was set to a delay of zero, the window extension was set to 0 µs, and the number of displayed events was set to 100. As needed, detector voltages (blue laser (488 nm) with B525 detector (525/50 nm) were modulated bring the signal within the linear range of the detector ($<10^5$). The delays were then manually changed with a step size of 1 µs until the detector readout from each laser was maximized. After optimizing the delay value, the delay window was reset to 2 µs, and the forward scatter threshold reduced to 19,000. Area scaling was then set such that the area and height measurements were equivalent, ensuring that the area, which was used in the assay was at least as large as the height. FSC and SSC voltages were adjusted using the 32 µm AccuCount beads to bring the beads above threshold and on-scale. Drop delay were optimized with the same 32 µm calibration beads to set a baseline. Basically, accudrop components consist of a diode laser mounted to the left of the sort block, a camera that views the images of the side stream and center stream, and an emission filter for viewing the fluorescence from calibration beads. The intensities of the streams are calculated during image processing as the delay is adjusted. With 32 (+/-2) um Sphero™ AccuCount ultra rainbow calibration beads, the drop delay value was adjusted in 0.03-drop increments until the left side stream intensity is greater than 95%. This drop delay established a baseline for subsequent manual drop delay calibration with the double emulsions to be sorted.

Double emulsions were resuspended in their outer solution without dilution. Forward and side scatter detector voltages were modulated as needed to keep the main population above threshold and on-scale for the SSC-A vs. FSC-A plot. The sorter was setup with the 130 µm nozzle and the break-off into single droplets was stabilized using adjustment of the frequency and amplitude and maintained at a gap of 12, per usual manufacturer recommendations. Sample pressure was maintained at 4 for all conditions. For sorting double emulsions in a library tube (without a corkscrew), 50-100 µL of double emulsions was transferred into a 1.2 mL polypropylene microtiter tube (Fisher Scientific, cat. # 02-681-376). The samples were manually resuspended by rotation before being loaded onto the cell sorter. During sorting, the samples were agitated 300 rpm rotation. For sorting double emulsions in a FACS tube (with a corkscrew), a 3D printed corkscrew was integrated onto the sample line of FACSAria II (BD) sample line following the sample line filter change procedure. Initially, the sample line was lowered to pass through the center hole of the corkscrew, positioning the extrusion part a few millimeters from the bottom of the corkscrew. The sample line was then raised back to its original position. An empty 5 mL FACS tube (Falcon, cat. # 352235) was subsequently installed to adjust the sample line height, ensuring a 2-3 millimeters gap between the sample line bottom and the FACS tube bottom. This adjusted height was maintained for subsequent sorting experiments. Then, approximately 600 uL of undiluted double emulsion suspension was placed into a FACS tube. These samples were also manually resuspended and then agitated at 300 rpm during sorting.

All samples were analyzed with manually set polygon gates for scattering events in the SSC-A vs. FSC-A plot and consecutive singleton gates in FSC-A vs. FSC-H and SSC-A vs. SSC-W plots. 10,000 events were recorded for each sample. Event rates ranged between 0-3000 Hz during sorting, and the sample pressure was maintained at 4.

Sorting was performed in purity mode. The corkscrew was attached to the sample line as described above. Droplet sorting delays were manually calibrated by sorting double emulsions onto a glass slide. At each delay step of 0.03 μs, 50 events detected by a scatter gate in the SSC-A vs. FSC-A plot were sorted onto a glass slide, and the number of recovered emulsions within the drops was counted under optical microscopy. Multiple steps were taken in each direction to establish a gradient. Steps were then taken in the positive direction until the number of recovered decreased. Additional measurements were then taken near the maximum, and the delay with the highest recovery rate was selected. 3-5 measurements of recovery rate were made at the maximum. Typical sort recovery was 75-80%. Then, approximately 60 uL of undiluted double emulsion suspension with 20 μM fluorescein in the inner aqueous phase and 600 uL of undiluted unlabeled double emulsion were placed into a 5mL FACS tube. A threshold gate for high fluorescein ($>10^3$) was created on the B525 histogram plot, and DEs within this gate were sorted into a 1.5 mL Eppendorf tube containing ~200 μL of outer buffer solution.
